# Molecular Mechanisms Underlying the Spectral Shift in Zebrafish Cone Opsins

**DOI:** 10.1101/2024.09.24.614827

**Authors:** L. América Chi, Shubham Kumar Pandey, Wojciech Kolodziejczyk, Peik Lund-Andersen, Jonathan E. Barnes, Karina Kapusta, Jagdish Suresh Patel

## Abstract

Visual pigments are essential for converting light into electrical signals during vision. Composed of an opsin protein and a retinal-based chromophore, pigments in vertebrate rods (Rh1) and cones (Rh2) have different spectral sensitivities, with distinct peak absorption wavelengths determined by the shape and composition of the chromophore binding pocket. Despite advances in understanding Rh1 pigments such as bovine rhodopsin, the molecular basis of spectral shifts in Rh2 cone opsins has been less studied, particularly the E122Q mutation, which accounts for about half of the observed spectral shift in these pigments. In this study, we employed molecular modeling and quantum mechanical techniques to investigate the molecular mechanisms behind the spectral difference in blue-shifted Rh2-1 (absorption peak = 467 nm, 122Q) and green-shifted Rh2-4 (absorption peak = 505 nm, 122E) zebrafish cone opsins. We modeled the pigments 3D structures based on their sequences and conducted all-atom molecular dynamics simulations totaling 2 microseconds. Distance analysis of the trajectories identified three key sites: E113, E181, and E122. The E122Q mutation, previously known, validates our findings, while E181 and E113 are newly identified contributors. Structural analysis revealed key features with differing values that explain the divergent spectral sensitivities of Rh2-1 and Rh2-4: 1) chromophore atom fluctuations and C5-C6 torsion angle, 2) binding pocket volume, 3) hydration patterns, and 4) E113-chromophore interaction stability. Quantum mechanics further confirms the critical role of residue E181 in Rh2-1 and E122 in Rh2-4 for their spectral behavior. Our study provides new insights into the molecular determinants of spectral shifts in cone opsins, and we anticipate that it will serve as a starting point for a broader understanding of the functional diversity of visual pigments.

## Introduction

Visual pigments are indispensable for an organism’s survival, playing a pivotal role in essential activities such as mating, foraging, and predator avoidance. Located in the rod and cone photoreceptors of the retina, these molecules are crucial for the initial conversion of light into electrical signals. Rod photoreceptors facilitate vision in dim light, while cone photoreceptors are responsible for daylight and color perception. Together, they enable the brain to interpret these signals as vision. Understanding the mechanisms underlying visual pigments’ function is not only fundamental but also central to diverse fields ranging from physiology to bioengineering.^1,2^

Each visual pigment is composed of two primary elements: an opsin protein and a retinal-based chromophore, with the chromophore attached to the protein via a protonated Schiff base (PSB) linkage. These pigments exhibit a remarkable diversity of spectral sensitivities, reflecting their adaptive responses to different environments. The range of spectral sensitivities for each visual pigment and the corresponding maximum absorption wavelengths (*λ_max_*) are influenced by the type of chromophore and the associated opsin protein. Given that the most common chromophore is 11-*cis* retinal,^3^ the shape and composition of the retinal binding pocket in the opsin protein are critical in determining the spectral sensitivity differences among visual pigments.^4–6^ For instance, an 11-*cis*-retinal molecule dissolved in an organic solvent like ethanol typically exhibits a *λ_max_*of approximately 440 nm.^7^ However, within visual pigments, this *λ_max_* can vary from about 420 to 560 nm, largely due to interactions with residues in the protein’s binding site. The shift in *λ_max_* from that of the PSB model compound in solution to that in the visual pigment is known as the “opsin shift”. This term is also employed to describe variations in *λ_max_* among different pigments. While progress has been made and some important insights have been gained, fully elucidating the molecular basis of spectral tuning in visual pigments remains a formidable challenge.

Vertebrate visual opsins are classified into five phylogenetic groups based on their absorption spectra: rod opsin or rhodopsin (RH1), rhodopsin-like cone opsins 2 (RH2), shortwavelength-sensitive cone opsins 1 and 2 (SWS1/SWS2), and medium/long-wavelength-sensitive cone opsins (M/LWS).^8^ Bovine rhodopsin (referred to as Rh1 from this point) was the first visual pigment with a determined experimental structure, leading to extensive research on its function and mechanisms.^9,10^ Recent advancements have also revealed the structure of human LWS cone opsin,^11^ broadening our understanding of visual pigments. However, experimental structures for other families, including RH2 cone opsin, are still lacking, leaving the spectral tuning mechanisms of RH2 cone pigments less explored. Fortunately, the high homology between RH1 and RH2 opsins allows the use of homology modeling to investigate the spectral tuning mechanisms of RH2 cone opsins. In particular, zebrafish, a model organism for understanding the mechanisms of vision and developing potential therapies for vision-related conditions, ^12^ possess four duplicated RH2 opsin genes (RH2-1 to RH2-4).^13^ The reconstituted photopigments from these genes (Rh2-1 to Rh2-4), with 11-*cis* retinal, exhibit a broad range of absorption spectra (450-540 nm), providing an excellent model to study spectral tuning mechanisms of RH2 in vertebrates.^13^

In general, at least 20 mutation sites in opsin pigments have been shown to cause notable spectral shifts via experimental reconstitution of vertebrate visual photopigments using a labor-intensive heterologous cell-culture system that measures absorption spectra.^14^ One substitution that causes a large spectral shift occurs at site 122. ^14,15^ Residue 122 is one of the 27 amino acids forming the retinal-binding pocket, located within 4.5 Å from 11-*cis* retinal, lining the *β*-ionone ring of the retinal chromophore in bovine rhodopsin (Rh1).^16^ Replacement of highly conserved glutamate by glutamine (E122Q) at this site has been specifically studied in Rh1, resulting in a spectral blue shift (shorter wavelengths) between 15 and 21 nm.^14^ One proposed mechanism in Rh1 suggests that the *β*-ionone ring of retinal is stabilized by rhodopsin’s electric dipole groups, particularly the electronegative carboxyl oxygen of GLU122, resulting in a green shift in the Schiff base’s excited state.^17^ On the other hand, the E122Q replacement resulted in a 13 nm blueshift to the coelacanth Rh2 pigment.^18^ In contrast, in zebrafish, the inverse amino acid substitution (Q122E) in Rh2-1 accounts for half of the spectral difference (19 nm out of 39 nm) between Rh2-1 and Rh2-4.^19^ Instead, the forward mutation (E122Q) in a named ancestor 1 explains almost half (15/32) of the spectral difference between ancestors 1 and ancestor 2. The same mutation in ancestor 3, explains the majority (14/18) of the spectral difference between ancestor 3 and Rh2-3. ^20^ Besides the remarkable importance shown for the E122Q/Q122E mutation, the molecular mechanism behind this mutation-induced spectral shift remains elusive.

In this study, we used molecular modeling and quantum mechanical techniques to explore the molecular mechanisms behind the spectral shifts in zebrafish Rh2-1 (*λ_max_* = 467 nm) and Rh2-4 (*λ_max_* = 505 nm) cone opsins, the most blue– and green-shifted Rh2 genes respectively, with special emphasis on the spectral shift induced by the E122Q mutation. We modeled the 3D structures of these pigments based on their opsin sequences and performed microscale atomistic classical molecular dynamics (MD) simulations, totaling 2 *µ*s, in a membrane and aqueous environment. From these simulations, we extracted both static and dynamic structure-based features to identify functionally critical amino acid sites, revealing both previously known and novel sites. We then proposed how these features contribute to understanding the molecular mechanisms underlying spectral shifts in these systems. Our key findings highlight the significant role of residues GLU113, GLU181, and GLN/GLU122 in the spectral shifts of Rh2-1 and Rh2-4 pigments.

## Results

### The global structural fluctuation differs between Rh2-1 and Rh2-4 pigments

While Rh2 pigments show similar flexibility patterns to Rh1, significant global structural changes have been observed between Rh2-1 and Rh2-4 photopigments. Both are 349 residue-long zebrafish proteins that have high sequence homology (83%) between them and with their bovine Rh1 paralog (66% and 71%, respectively) as shown in Figure S1 (see Supplemental Material). The structure of Rh2-4 photopigment highlighting characteristics of an opsin protein is shown in Figure 1a-b, including the seven transmembrane helices, the antiparallel sheet motif interacting with the chromophore from the extracellular side, and the 11-*cis* retinal chromophore attached to residue LYS296 in helix 7. We first analyzed the root mean square deviation (RMSD) of the C*α* atoms for both Rh2-1 and Rh2-4 pigments to observe global fluctuations. We observed that the systems reached equilibrium within the first nanoseconds of the MD simulation and maintained stable structures throughout (Figure S2 of the Supplemental Material). Figure 1c-d shows the root mean square fluctuation (RMSF) of the C*α* atoms for the Rh2-1 and Rh2-4 and the corresponding RMSF values are mapped onto their 3D structures. Experimental *β*-factors for Rh1 are shown for comparison. Overall, high fluctuations occur in the coil/loop/*β*-sheet inter-helical regions in all systems. The Rh2-4 RMSF profile shows significantly higher fluctuations at the N-terminus, in coil/loop/*β*-sheet inter-helical regions, and the first segment of helix 6.

**Figure 1:**
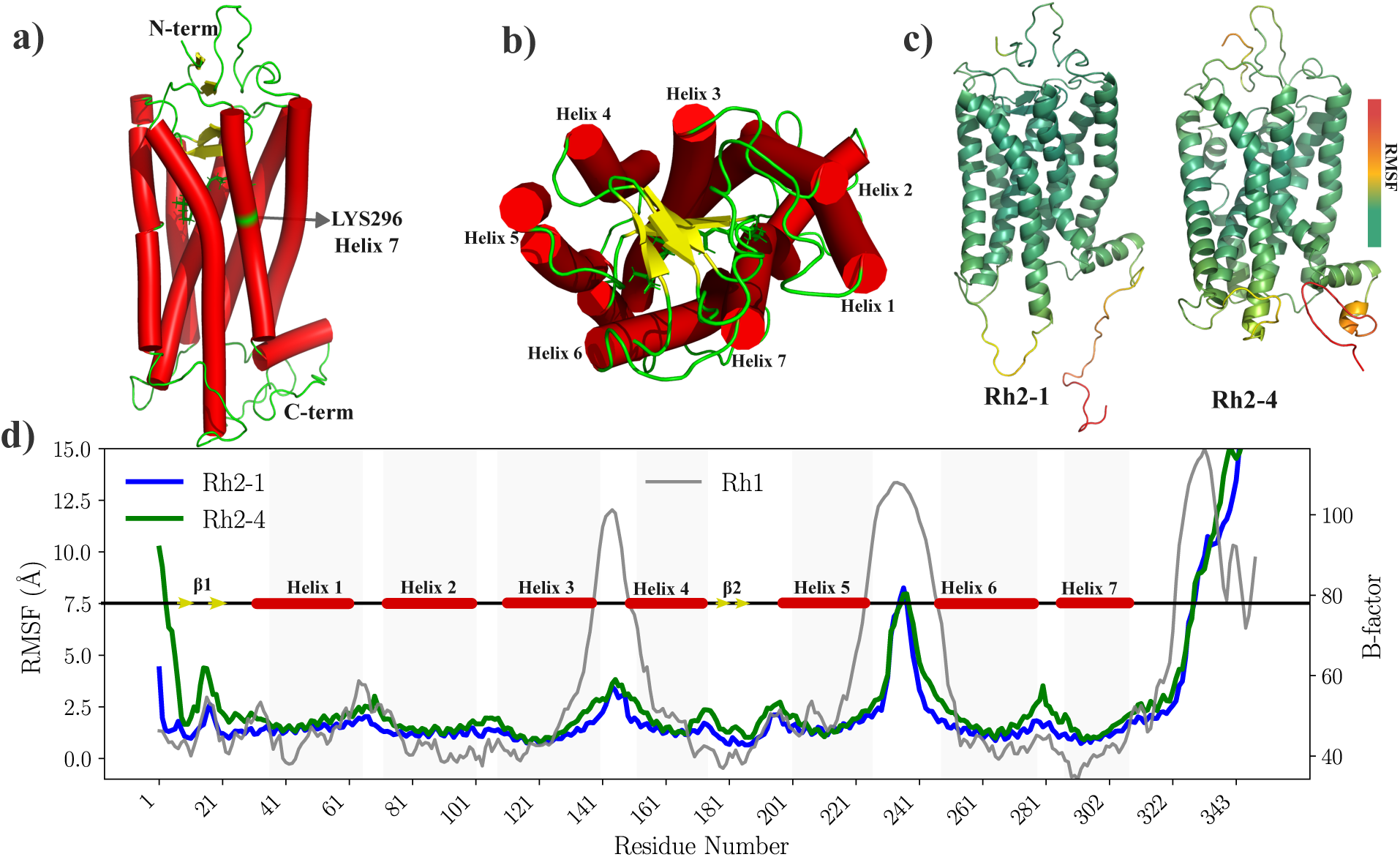
Comparison of structural fluctuations in Rh2-1 and Rh2-4 pigments. The Rh2-4 pigment is shown in side (a) and top (b) views, with the seven transmembrane helices depicted in the tube representation, the chromophore in the stick representation, and the antiparallel sheets motif in the cartoon representation. Root Mean Square Fluctuation (RMSF) profiles of Rh2-1 (blue line) and Rh2-4 (green line) visual photopigments during 1*µ*s long MD simulations are mapped onto 3D structures (c) and plotted along amino acid residue numbers (d). Experimental *β*-factors for bovine rhodopsin (Rh1) are shown in the gray line as a reference (Right Y axis).^10^ Shadow spots highlight the seven transmembrane helices reported for Rh1.

### The local environment diverge in Rh2-1 and Rh2-4 pigments

Local structural changes were observed between Rh2-1 and Rh2-4 pigments. The volume of the chromophore pocket over time (Figure 2a) shows differences, with Rh2-4 maintaining a larger pocket volume compared to Rh2-1. The residues surrounding the chromophore pocket are presented in Figure 2b, showing different environments in the Rh2-1 and Rh2-4 systems. In general, the chromophore is surrounded by charged (GLU) and polar (SER) residues near the protonated nitrogen or Schiff base proton (N^+^/SBH^+^) region and hydrophobic (PHE, TRP, ALA, LEU) residues around the *β*-ionone ring. In both systems, GLU113 and SER186 line the SBH^+^. In addition, Rh2-1 chromophore is lined by GLU181, SER94, PHE91, and CYS187, while Rh2-4 is lined by ALA117. For the *β*-ionone ring, both systems have LEU125 and TRP265 as lining residues. However, Rh2-1 has additional residues PHE261, PHE212, GLN122, and ALA269, while Rh2-4 has GLU122, GLY121, and HIS211. The chromophore in Rh2-1 and Rh2-4 is surrounded by residues that are also found in the binding pocket of bovine Rh1, as shown in Figure S3 in the supplemental material. An important difference is that of the 27 sites found in the Rh1 pocket, Rh2-1 coincides with 25 and Rh2-4 with 23. Among the residues shared with Rh1, variations are noted at several sites: site 94 hosts a threonine in Rh1 and Rh2-4, but a serine in Rh2-1; site 122 hosts glutamate in Rh1 and Rh2-4, and glutamine in Rh2-1; site 189 has isoleucine in Rh1, but is replaced by proline in both Rh2-1 and Rh2-4, and site 295 contains an alanine in Rh1, but a serine in both Rh2-1 and Rh2-4. The presence or absence of specific residues near the chromophore may influence how opsins fine-tune their absorption properties. To assess whether these changes are consistent on average, we performed a detailed distance analysis, as discussed in the following section.

**Figure 2:**
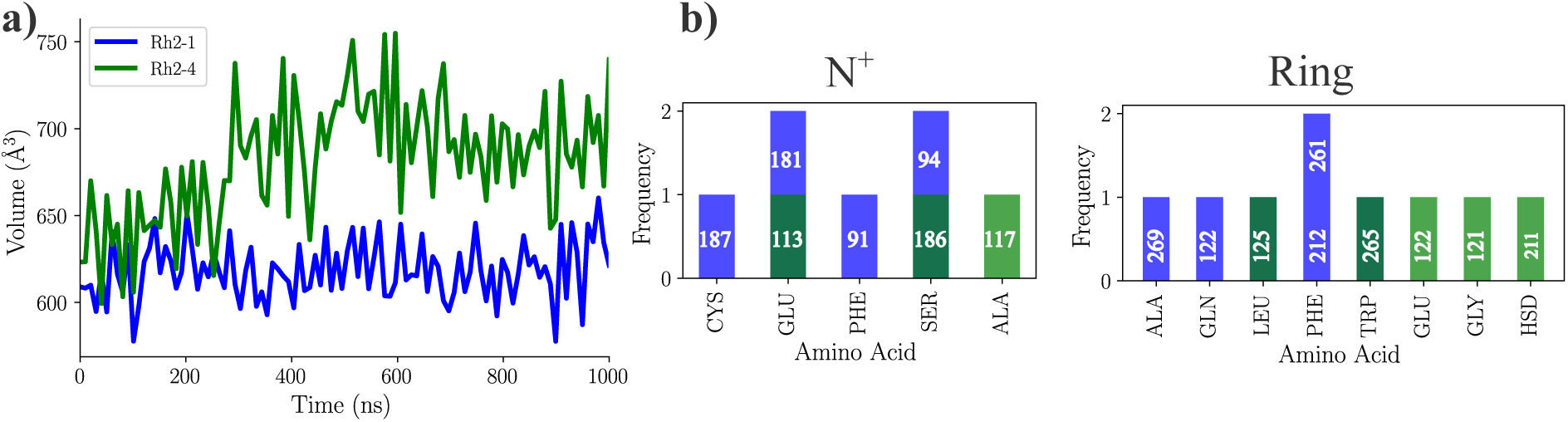
Variations in the local structure around the chromophore in Rh2-1 and Rh2-4 pigments. a) The volume of the chromophore pocket over time. b) Residues lining the chromophore pocket (*β*-ionone ring and N^+^) for representative structures.

### Distance analysis pinpoint potential critical sites

Notable changes in the distances between the centers of mass of helix 7, which contains the chromophore, and the *β*_2_-sheet motif or helix 3 were observed in Rh2-4 compared to Rh2-1 (Figure S4 in the Supplemental Material). While all helices exhibit some degree of bending, Helix-5 shows significantly more bending in Rh2-1 compared to Rh2-4 (Figure S5 in the Supplemental Material). To examine more closely which sites might be involved in significant structural changes, we measured the relative differences in distances between the residues and the chromophore (Figure 3, revealing that several residues in Rh2-4 tend to be further/closer from the chromophore than those in Rh2-1. Residues at sites 181, 292, and 187 were significantly farther from the N^+^, whereas residues at sites 90, 113, and 117 remained closer. Residues at sites 122, 211, and 212 were significantly farther from the N^+^, whereas residues at site 268 remained closer. From the sites identified in the distance analysis, we chose to further investigate sites 113, 122, and 181, which have been moderately to extensively discussed in the bovine rhodopsin literature.

**Figure 3:**
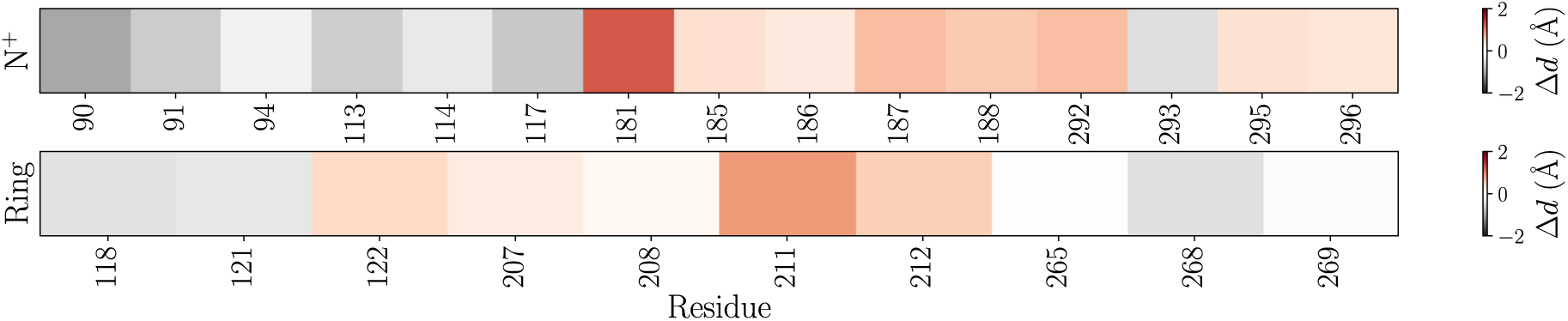
Relative difference in the distances of the residues from the chromophore. This was measured specifically for those residues whose C*α* atoms were within 8 Å of the chromophore. We first measured the distance between C*_α_* atoms and the protonated nitrogen/*β*-ionone ring, then averaged these distances across the entire simulation. Thus, we compared the difference between the same residue-chromophore pair in Rh2-4 to those in Rh2-1. In this context, grey indicates that the average distance from a specific residue to the chromophore domain (protonated nitrogen N^+^ or *β*-ionone ring) is shorter in Rh2-1 than in Rh2-4, while orange signifies that the distances are greater.

### Chromophore structure dynamics vary within Rh2-1 and Rh2-4 proteins

Chromophore dynamics vary within Rh2-1 and Rh2-4 proteins. Figure 4a displays the chemical structure of the chromophore covalently attached to LYS296 (LYR). The RMSF of the LYR C*α* atoms is shown in Figure 4b, where Rh2-4 exhibits higher fluctuations compared to Rh2-1, especially towards the *β*-ionone ring. In Figure 4c, the LYR radius of gyration (R*_g_*) over time shows that Rh2-4 maintains a slightly larger radius, reflecting a more extended chromophore conformation. Figure 4d-e presents the key angle distributions. The peak in the histograms for the CD-CE-N^+^-C15 torsion angle in Rh2-1 is noticeably shifted to higher angles compared to Rh2-4. The histogram for the C7-C6-C1-C16 torsion angle in Rh2-1 is monomodal, whereas in Rh2-4 it is bimodal. The peak in Rh2-1 overlaps with the higher angle peak in Rh2-4. The histogram for the C5-C6-C7-C8 torsion angle in Rh2-1 is monomodal, whereas in Rh2-4 it is bimodal. The peak in Rh2-1 overlaps with the negative angle peak in Rh2-4. The peaks in the histograms for the CA-N^+^-C13/C14-C10-C2 geometric angles in Rh2-1 are noticeably shifted to lower angles compared to Rh2-4.

**Figure 4:**
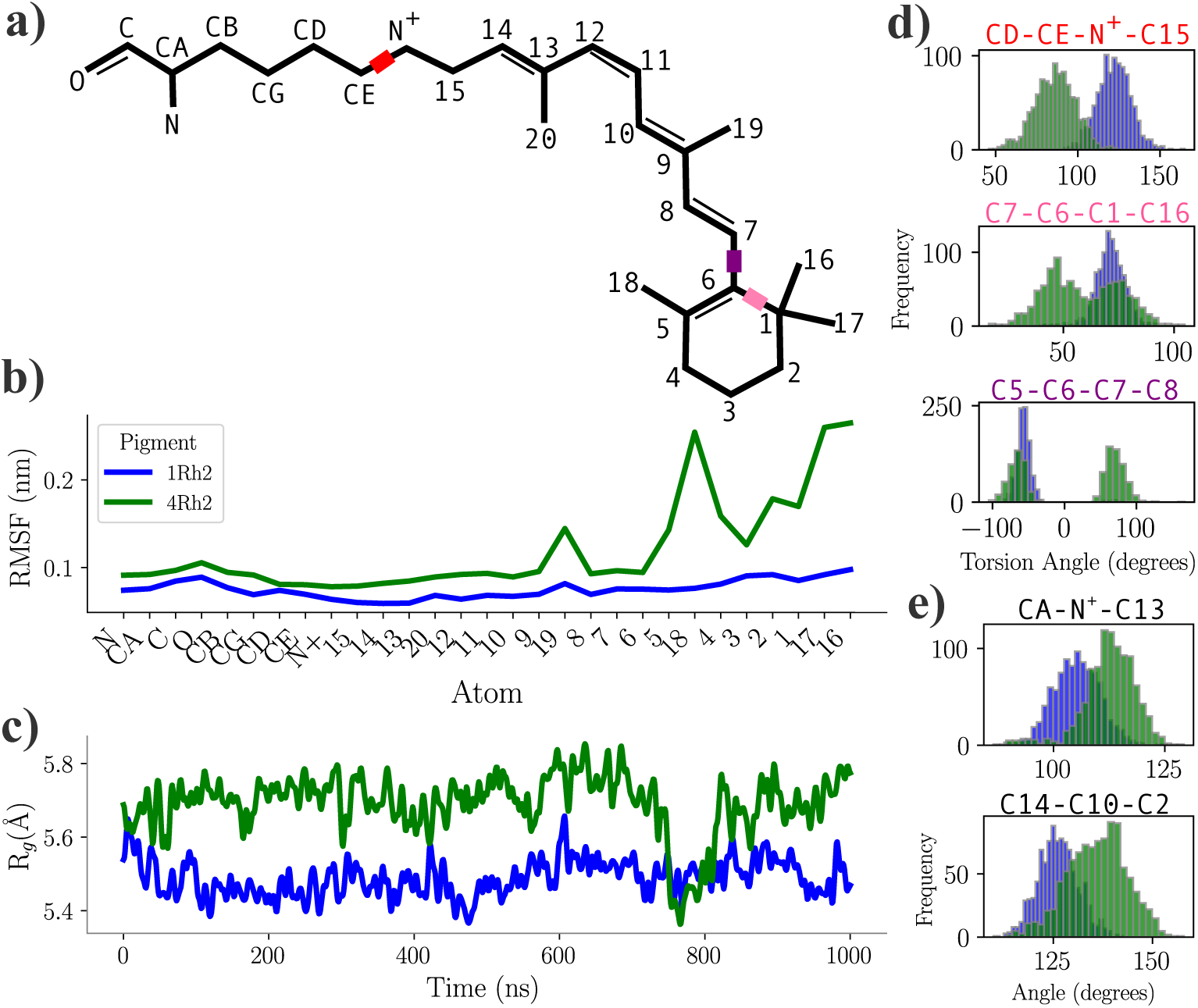
Chromophore dynamics and structural analysis within Rh2-1 and Rh2-4 proteins. a) The chemical structure of the chromophore plus LYS296 (LYR), with specific atoms and torsion angles analyzed. b) The RMSF of the LYR atoms in Rh2-1 (blue) and Rh2-4 (green) proteins. c) The radius of gyration (R*_g_*) for residue LYR over time. d) Torsion angle distributions for key bonds within the chromophore: CD-CE-N^+^-C15, C7-C6-C1-C16, C5-C6-C7-C8. e) Geometric angle distributions for CA-N^+^-C13, and C14-C10-C2.

### Chromophore interactions with its environment in Rh2-1 and Rh2-4

Chromophore-opsin interactions show differences between Rh2-1 and Rh2-4 pigments. Figure 5 show a detailed view of the interactions of the residue with LYR. Hydrogen bonds (HBs) are observed between the SBH^+^ and the GLU113 counterion in both systems. In the Rh2-1 system, additional HBs are formed between the NH of LYS296 and ALA292, and between the CO of LYS296 and VAL300. Conversely, in the Rh2-4 system, an HB is formed between the NH of LYS296 and PHE293. The additional presence of GLU181 in the Rh2-1 system is noteworthy; GLU181 is oriented toward the chromophore, creating a differentially positively charged environment around the chromophore, which is not observed in the Rh2-4 system. GLN122 polar residue is oriented towards the *β*-ionone ring of the chromophore in Rh2-1, whereas the positively charged residue GLU122 is directed away from the chromophore in Rh2-4. Additionally, the chromophore pocket is more hydrated, and lipid molecules stay closer in Rh2-4 compared to Rh2-1 (Figures S6-S7 from Supplemental Material).

**Figure 5:**
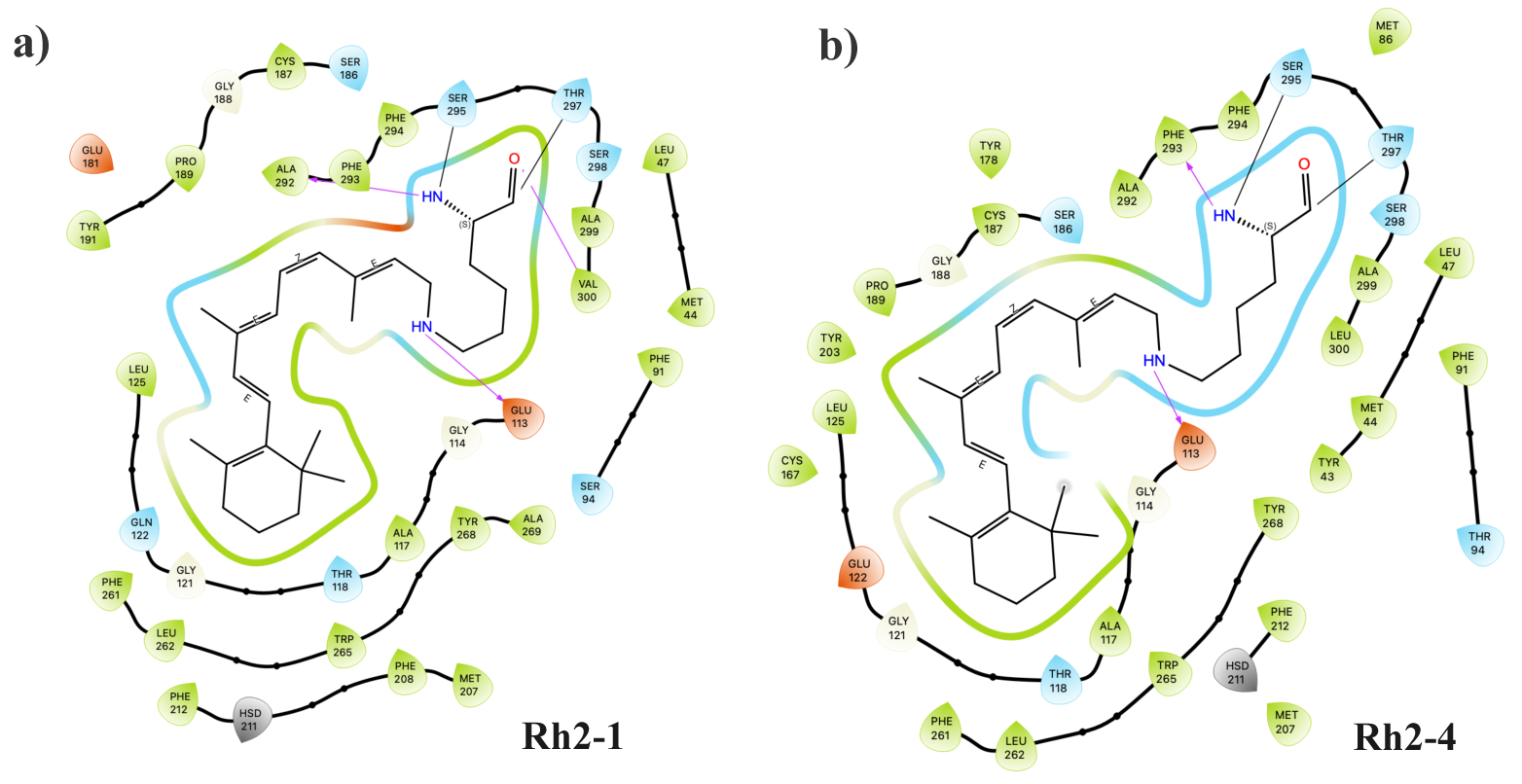
Chromophore-opsin interactions in Rh2-1 and Rh2-4 pigments. The two-dimensional schematics provide detailed views of residue interactions, including hydrogen bonds (magenta arrows), hydrophobic (green), polar (blue), and charged positive (red). The diagram was built with the Maestro Schrödinger suite.^21^

**Figure 6:**
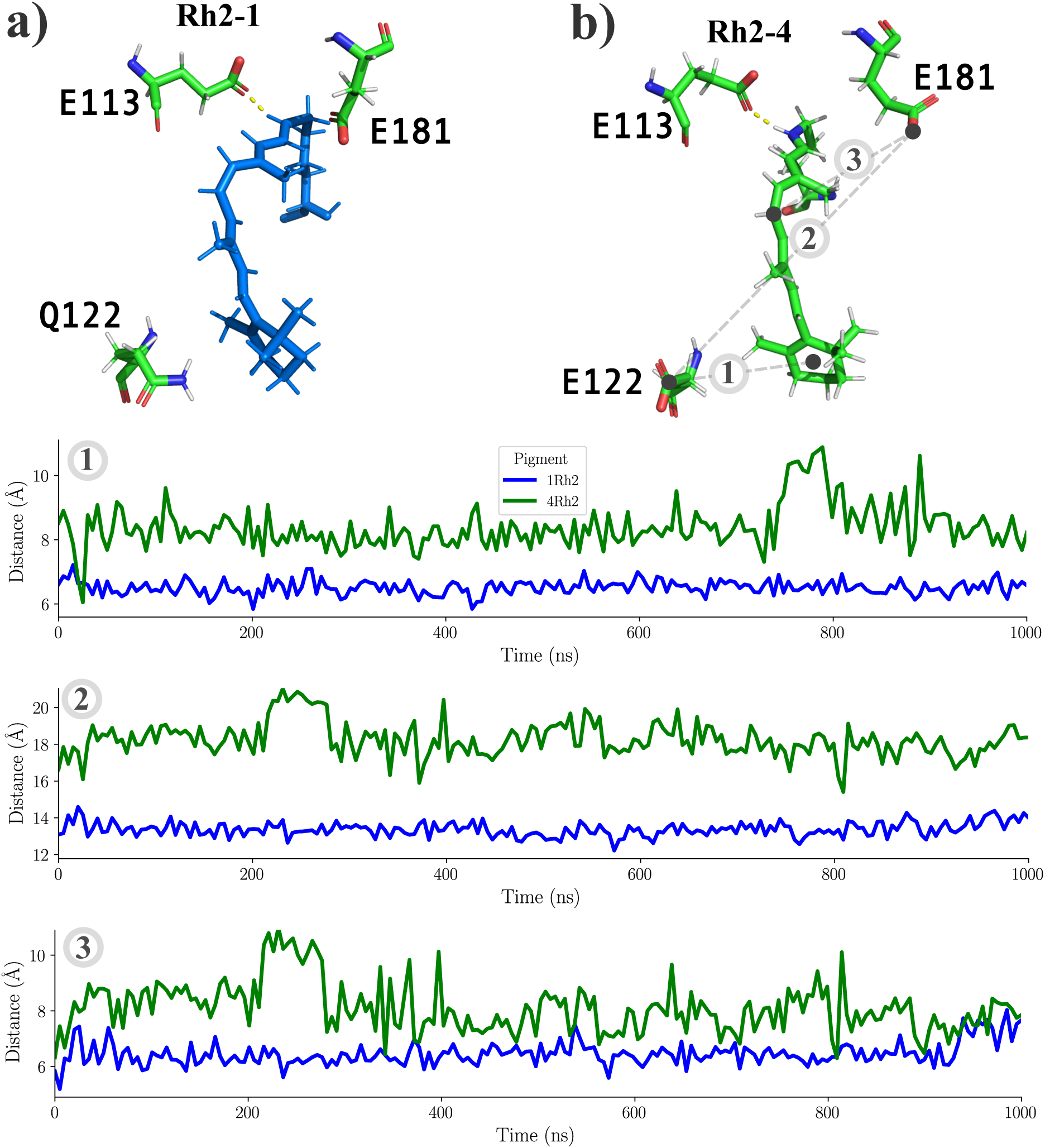
Distance and orientation between key residues in. a) Rh2-1 or b) Rh2-4. 1) The distance between the GLU/GLN122 residue and the *β*-ionone ring of the chromophore. 2) The distance between the GLU181 residue and the polyene region of the chromophore. 3) The distance between the GLU181 residue and residue GLU/GLN122.

### The key role of GLU113, GLU181 and GLU/GLN122 in differential pigments behavior

The chromophore is significantly influenced by key residues E113, E181, and GLU/GLN122. Figure 8 shows the relative positions and orientations of these residues to the chromophore or between them in both systems. Figure 8a depicts the orientation of each residue around LYR of Rh2-1, while Figure 8b represents Rh2-4 case. The distance between the GLU/GLN122 residue and the *β*-ionone ring of the chromophore is larger in the Rh2-4 system. The distance between the GLU181 residue and the polyene region of the chromophore is larger in the Rh2-4 system. The distance between the GLU181 residue and residue GLU/GLN122 is larger in the Rh2-4 system.

Residue GLU113, the counter ion for the PBS^+^, shows a differential behavior in Rh2-1 compared to Rh2-4. In Rh2-1, GLU113 can form HBs with NH^+^ of the SBH^+^, OH of SER94, or OH from water molecules (Figures S8-S9 from Supplemental Material). While the interaction with NH^+^ occurs frequently during the simulation, with an occupancy of 99%, the interaction with SER94 is less frequent, with an occupancy of 80%, and even lower for the water molecules, with an occupancy of 18%. In Rh2-4, GLU113 can form HBs with NH^+^ of the SBH^+^, OH of SER186, or OH from water molecules. The interaction with NH^+^ occurs frequently during the simulation, with an occupancy of 92%, the interaction with SER186 with an occupancy of 95%, and lower for the water molecules, with an occupancy of 19%. We also looked at the distances between the oxygen atoms of the GLU113 counterion and the N+ and angles between the donor-acceptor (Figure S8 from the Supplemental Material), and we found that the distances and angles are more frequently within the ideal range for hydrogen bond formation in Rh2-1 compared to Rh2-4. In summary, GLU133-SBH^+^ interaction seems more stable in Rh2-1 than in Rh2-4.

Residue 122, the residue whose change Q/E accounts for an important shift in the peak spectral absorption, shows a differential behavior in Rh2-1 compared to Rh2-4. In Rh2-1, GLN122 can form HBs with CO backbone atoms from H211 or NH-backbone/NH side chain atoms from TRP126 (Figures S10-S11 from Supplemental Material). The interaction with H211 occurs with an occupancy of 93%, and the interaction with TRP126 is less frequent, with an occupancy of 80%. In Rh2-4, GLU122 can form HBs with NH-backbone/NH side chain atoms from TRP126 or NH backbone atoms from CYS167. The interaction with TRP126 with an occupancy of 2.7%, and even lower for CYS167, with an occupancy of 0.4%. In summary, residue 122 has a more stable HB network in Rh2-1 than in Rh2-4.

Residue GLU181 exhibits differential behavior in Rh2-1 compared to Rh2-4. In Rh2-1, GLU181 can form HBs with the OH side-chain groups from TYR268 and SER186 or OH from water molecules (Figures S9-S10 in the Supplemental Material). The interaction with TYR268 has an occupancy of 94%, the interaction with SER186 has an occupancy of 93% and it interacts with 1 water molecule with an occupancy of 13%. In Rh2-4, GLU181 can form HBs with OH side-chain groups from TYR268, TYR191/TYR192, or water molecules. The interaction with TYR268 has an occupancy of 99.5%, interactions with TYR191 and TYR192 have lower occupancies of 41% and 39%, respectively and it interacts with 10 water molecules with occupancies below 25%. In summary, residue GLU181 has a more stable HB network with other residues in Rh2-1 than in Rh2-4.

### Electronic effects of opsin shifts in Rh2 photopigments through QM/MM optimization and TDDFT analysis

To investigate the electronic effects influencing the experimentally observed opsin shift, we conducted QM/MM (quantum mechanics/molecular mechanics) optimization following quantum mechanics excited state calculations. For the initial QM/MM optimization, the chromophore and GLU113 residues were included in the QM region as entire residues (including a side chain and a backbone). Analysis of the optimized structure (Figure 7a) revealed that, in addition to forming a salt bridge and hydrogen bond with GLU113, the NH^+^ of the SBH^+^ engaged in an additional *π*-cation interaction with PHE293 in the Rh2-1 protein. This interaction was absent in the Rh2-4. The hydrogen bonds between the LYR296 backbone and ALA292 and VAL300 remained unchanged for Rh2-1, as did the hydrogen bond between the chromophore backbone and PHE293 for Rh2-4. A newly formed *π*-cation interaction suggested that including PHE293 in further optimizations could improve accuracy. The preliminarily optimized structure was then re-optimized, with LYR as an entire residue and GLU113 and PHE293 as side chains included in the QM region. Despite this adjustment, the 3D structures of both Rh2 photopigments remained mostly unchanged (Figure 7b), except for the hydrogen bond with ALA292 being broken for Rh2-1. To assess chromophore-opsin’s absorption properties, LYR interactions were subsequently analyzed to select specific residues for TDDFT calculations. Various functional/basis set combinations, both in the gas phase and water solvent, produced consistent results, with the absorption maximum for Rh2-1 showing a significant redshift compared to Rh2-4 (23–37 nm, method-dependent). However, this trend contradicted known experimental data, indicating that the optimized structure was insufficient to describe the Rh2 system accurately. Nonetheless, the TDDFT results (Table S1 in Supplemental Materials) allowed us to benchmark the most effective method combinations regarding their quality and computational time.

**Figure 7:**
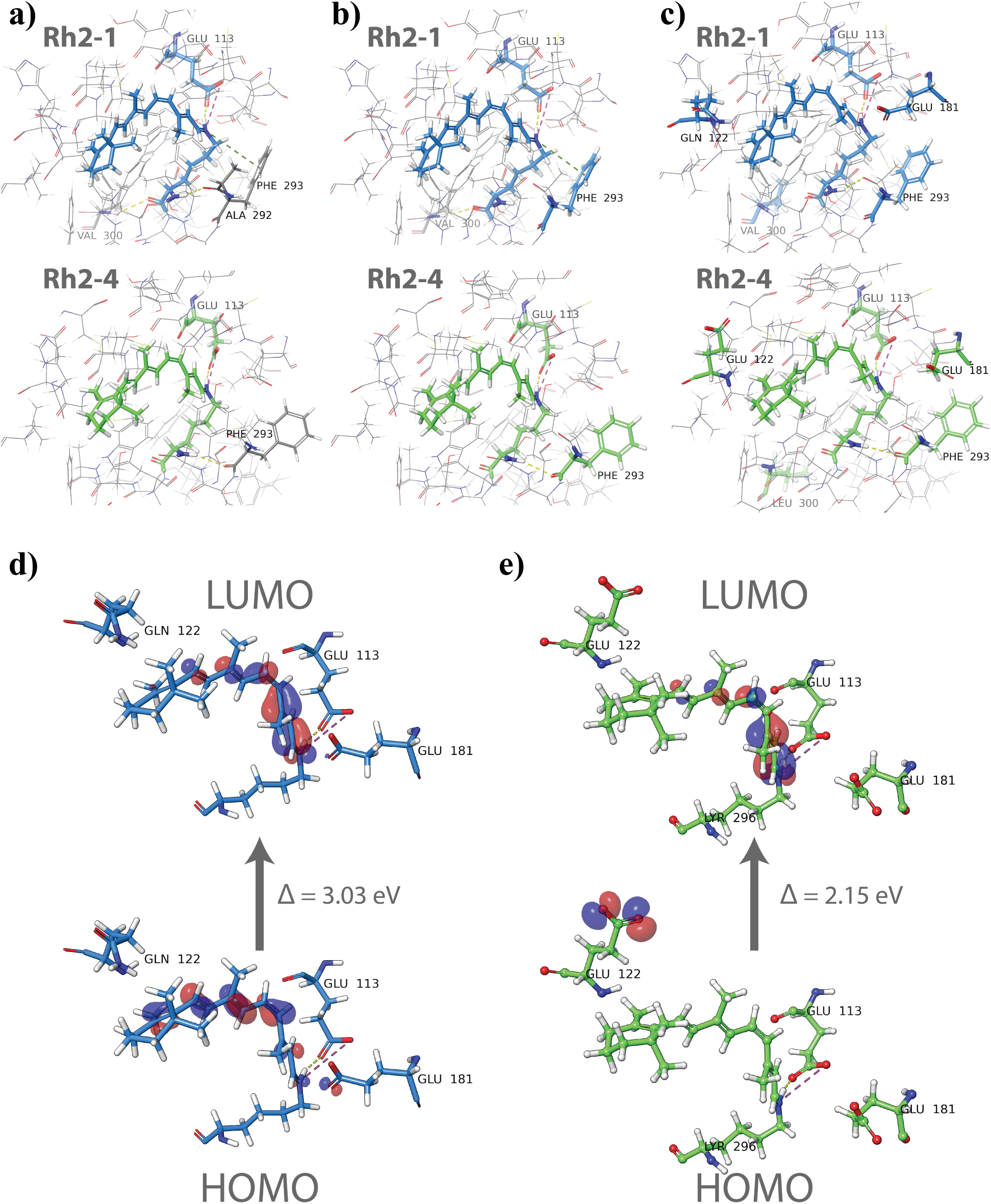
Results of the Rh2 proteins’ QM/MM optimization. Geometries of Rh2 with (a) LYR296 and GLU113 as entire residues, (b) LYR296 as an entire residue and GLU113 and PHE293 as side chains, and (c) LYR296 as an entire residue and GLU113, GLN/GLU122, GLU181, PHE293, and VAL/LEU300 as side chains. QM region is colored in blue or green, while the MM region is colored by element. Visualization of the highest occupied and lowest unoccupied molecular orbitals for (d) Rh2-1 and (e) Rh2-4 photopigments.

To achieve a more accurate QM/MM optimization, the second attempt included additional residues in the QM region: LYR296 (entire residue), GLU113, GLU/GLN122, GLU181, PHE293, and LEU/VAL300 as side chains. This refinement eliminated the *π*-cation interaction in the optimized structure of Rh2-1 (Figure 7c). The newly optimized Rh2-1 structure also featured a hydrogen bond between the chromophore and ALA292 (as in the original structure) but lacked one with VAL300. The Rh2-4 structure exhibited only minor changes. HOMO and LUMO orbitals were visualized for optimized structures (Figure 7d–e). The LUMO was similar in both Rh2 systems, spanning the chromophore between the NH^+^ of the SBH^+^ and *β*-ionone ring. However, the HOMO of Rh2-1 had significantly lower energy compared to Rh2-4, being shifted toward the *β*-ionone ring and partially located on GLU181. In contrast, the HOMO of Rh2-4 was solely situated on GLU122. The HOMO-LUMO gap was larger for Rh2-1 (3.03 eV) compared to Rh2-4 (2.15 eV). These findings supported the experimental results, indicating a red (bathochromic) shift in the absorption *λ*_max_ for Rh2-4. TDDFT calculations were performed on the newly optimized pigment structures to confirm this further.

The QM region of the optimized structure was extracted and used for single-point and TDDFT calculations, which were performed using two different method combinations based on the benchmark results shown in Table S1 (Supplemental Materials): the HF/3-21G level of theory in the gas phase and the CAM-B3LYP/6-31G_JSKE level of theory in water solvent (using the conductor-like polarizable continuum model). Both methods showed a bathochromic shift for the Rh2-4 protein, aligning with experimental data. The Hartree-Fock (HF) method with the economical, small split-valence basis set 3-21G in the gas phase revealed a more substantial *λ*_max_ difference of 15 nm between Rh2-1 and Rh2-4, with absorption peaks of large intensity, as shown in Figure 8a. In contrast, using the CAM-B3LYP hybrid exchange-correlation functional with our previously developed, physically justified augmented 6-31G_JSKE basis set^22^ resulted in less intense peaks and a smaller *λ*_max_ difference of 5 nm (Figure 8b).

To further explore these observations, HOMO and LUMO orbitals were visualized (Figure 8c-d). For the Rh2-4 pigment, similar orbital shapes were observed with both methods, yielding HOMO-LUMO gap values of 6.36 eV with HF/3-21G and 5.26 eV with CAM-B3LYP/6-31G_JSKE. However, a significant difference was observed in the Rh2-1 pigment. At the CAM-B3LYP/6-31G_JSKE level, the HOMO location resembled the distribution obtained from the QM/MM optimization with B3LYP/cc-pVTZ(-f) (Figure 7d), predominantly located between the *β*-ionone ring and the NH^+^ group of the chromophore, with partial distribution on GLU181. In contrast, the HF/3-21G calculation showed the HOMO exclusively on GLU181. These differences led to substantial variations in the calculated HOMO-LUMO gaps, with HF/3-21G yielding a gap of 8.24 eV, while CAM-B3LYP/6-31G_JSKE produced a significantly lower value of 5.37 eV. The larger gap difference observed with Hartree-Fock was consistent with the more pronounced shift in the absorption maximum, whereas the smaller HOMO-LUMO gap obtained with CAM-B3LYP corresponded to a smaller *λ*_max_ difference. Detailed results of these calculations are presented in Table S2 (Supplemental Materials).

**Figure 8:**
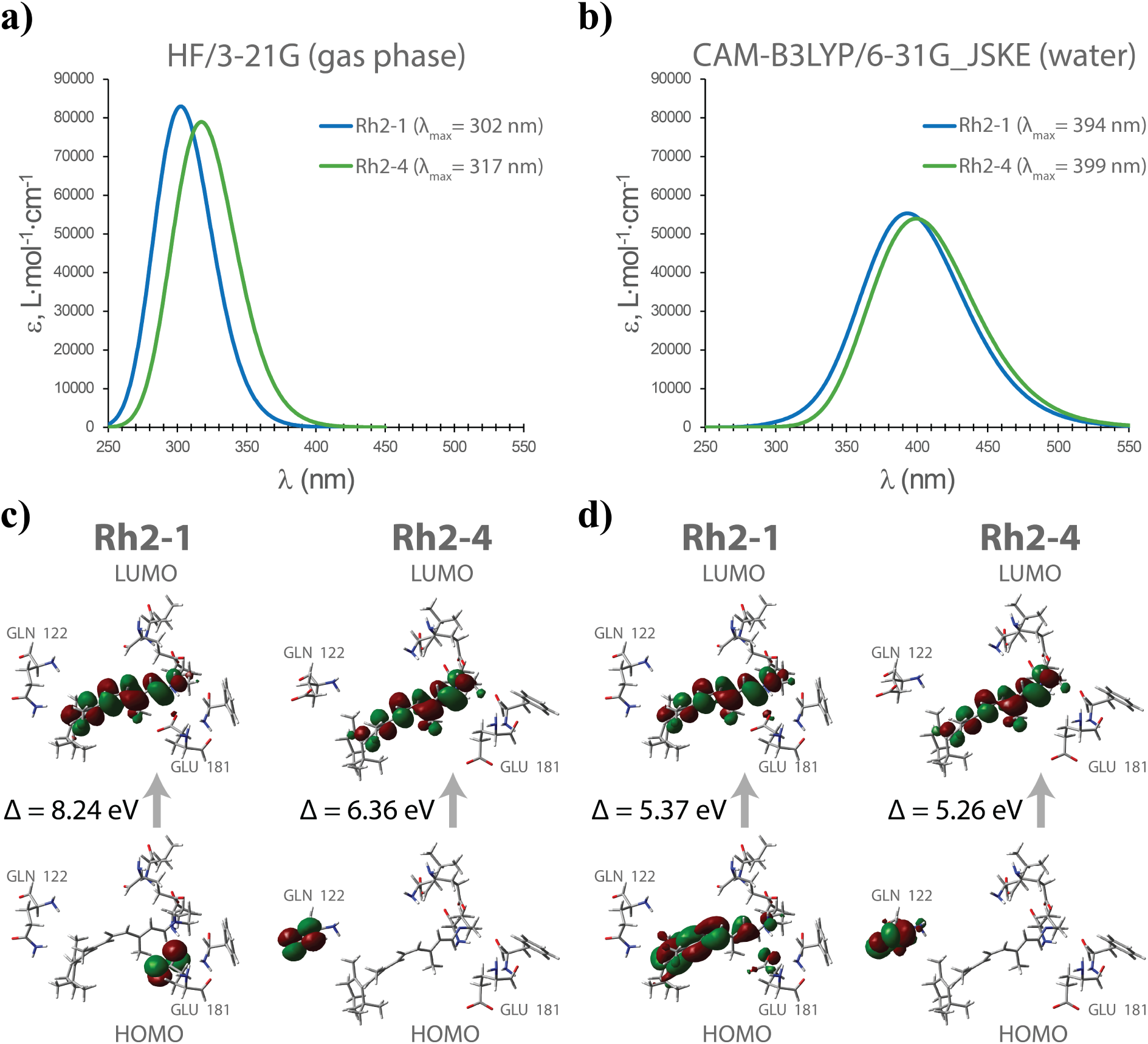
Absorption properties of Rh2 photopigments. UV-Vis spectra calculated using (a) the HF/3-21G level of theory in the gas phase and (b) the CAM-B3LYP/6-31G_JSKE level of theory in water solvent. Visualization of HOMO and LUMO performed using (c) the HF/3-21G level of theory in the gas phase and (d) the CAM-B3LYP/6-31G_JSKE level of theory in water solvent.

## Discussion

In this study, we used molecular modeling techniques to gain insight into the largely unexplored molecular mechanisms behind the spectral shifts of Rh2 cone opsins, in particular the zebrafish Rh2-1 and Rh2-4. Taking advantage of their homology to bovine rhodopsin, we constructed 3D models of these pigments and subsequently performed microscale classical all-atom MD simulations on the systems. From these simulations, we extracted both static and dynamic structure-based features through a top-down structural analysis, identifying functionally critical amino acid sites, including both previously known and novel ones. We performed a detailed analysis of the characteristic structural features of these sites and proposed how they may contribute to the molecular mechanisms driving spectral shifts in these systems.

Our study leveraged the significant homology of zebrafish Rh2-1 and Rh2-4 cone opsins to bovine Rh1 rhodopsin to construct models in the absence of experimental structures. The careful selection of optimal models, together with the rapid achievement of equilibrium and well-preserved structures observed during extensive MD simulations, ensures the reliability of our approach and the insights gained from our MD simulations. Furthermore, we anticipated some degree of structural dynamic similarity between Rh1, Rh2-1, and Rh2-4 due to substantial sequence homology. We confirmed that the global flexibility patterns of Rh2-1 and Rh2-4 closely resemble those observed in bovine rhodopsin, especially in interhelical loop regions and terminal domains, i.e., regions with higher RMSF values were consistent with elevated experimental *β*-factor values seen in Rh1 crystal structures^10^ and similar computational studies.^23,24^ A key difference is that, unlike Rh2-1, Rh2-4 exhibits a greater overall degree of fluctuation, particularly at the base of helix 6. In Rh1, spin-label mobility studies have indicated significant shifts at the cytoplasmic end of this helix, often associated with G-protein binding during activation.^10,25^ This observation highlights the need for further studies on Rh2 systems in this regard. In particular, no other computational studies have yet investigated the dynamics of Rh2 opsins in zebrafish, which provides a strong motivation for our research, but also makes it difficult to compare our results with the existing literature. In general, the retinal binding pocket volume in Rh2-1 and Rh2-4 zebrafish cone opsins are notably larger, approximately double that of bovine rhodopsin Rh1,^26^ suggesting a much tighter packing within rhodopsin’s chromophore binding pocket compared to cone opsins.^19^

The larger chromophore binding pocket in Rh2-4 compared to Rh2-1 reflects a differential microenvironment surrounding the chromophore. This result goes in line with the observation that Rh2-4 reacted more quickly in the presence of hydroxylamine than Rh2-1, suggesting the possibility of differences in the accessibility of the chromophore binding pocket between these two opsins.^19^ The variation in pocket shape observed in Rh2 cone opsins may allow adaptation to subtle differences in chromophore size, flexibility, and torsion angle. Alternatively, the microenvironment within the pocket could influence the structure of the chromophore and contribute to the differences observed between the two systems. Particularly, the dihedral angle around the C6-C7 single bond exhibited differential distribution in Rh2-1 and Rh2-4. In rhodopsin, this torsional angle is a well-known factor contributing to spectral shifts, where absorption energy increases with greater torsion. ^27–30^ These structural adaptations are likely to fine-tune the absorption properties of the opsins by influencing how the retinal fits and interacts within the pocket. When the binding pocket is larger and more flexible, the chromophore can interact with surrounding residues more freely, which may change its electronic environment and make it easier for the chromophore to transition to its excited state in Rh2-4 than in Rh2-1. In terms of composition, the chromophore in Rh2-1 and Rh2-4 is surrounded by residues that are also found in the binding pocket of bovine Rh1. Although many sites/residues are conserved across the Rh1, Rh2-1, and Rh2-4 binding pockets, key differences between Rh2-1 and Rh2-4 could contribute to functional variations between these visual pigments. For instance, site 122, which contains glutamate in Rh1 and Rh2-4 but glutamine in Rh2-1, has been extensively studied in the Rh1 system but has received comparatively less attention in the Rh2 systems. In the other hand, site 94, which contains threonine in Rh1 and Rh2-4 but serine in Rh2-1, warrants further investigation due to its potential impact on pigment function.

It is widely recognized that the interaction between the chromophore and the surrounding protein environment plays a crucial role in altering the absorption maxima of opsin pigments.^31,32^ Residues, positioned strategically near the chromophore, can differentially influence the chromophore’s electronic environment, thereby altering its absorption spectrum. Consequently, measuring the differential distance of binding site residues from chromophore main domains emerged as a logical approach. We first examined the spatial relationships between various structural domains relative to Helix 4 which contains the chromophore. We identified significant positional shifts in Helix 3 and the *β*_2_ segment. Furthermore, using a distance analysis we were able to identify functionally critical amino acid sites. Particularly residues 113, 122, and 181, experienced differential distance distribution from the chromophore and thus might play pivotal roles in the color tuning of Rh2 zebrafish cone opsins as these amino acids involve chemical properties that could alter the dielectric and electrostatic character of the chromophore-binding pocket. Residue 122 has been pointed out in previous mutagenesis analysis as responsible for a significant part of the Rh2 and Rh1 spectral shift.^14,18–20^ Residue 113 has mostly been studied on Rh1, mainly as important for activation^33^ but also as has been mentioned as a contributor to spectral shift. On the other hand, residue 181 has been related to Rh1 activation,^34^ a few studies claim that it does not contribute significantly to the Rh1 spectral tuning^35^ while others suggest the contrary.^36^ Such findings underscore the utility of distance analysis as a tool not only for structural but also for functional annotation in photoreceptive proteins. The simplicity of the distance analysis, combined with its ability to yield insights into functionally significant molecular interactions, provides a robust framework for future studies. This approach can be particularly useful in cases where experimental data are scarce or when rapid screening of potential functional sites is required.

Multiple factors typically contribute to the mechanisms underlying spectral shifts in opsins. Some of the most common factors include 1) changes in the strength of interactions of the electrostatic interaction between the PSB group of the chromophore and its counterion;^37,38^ 2) placement of full or partial charges along the retinal;^4–6^ 3) an alteration in the polarity or polarizability of the environment of the chromophore-binding site; ^39^ and 4) twisting along bonds of the polyene chain.^7^ Our results suggest that the opsin shift observed in Rh2-1 and Rh2-4 zebrafish cone opsins might involve a combination of changes involving these factors and critical residues at sites 113, 122, and 181.

GLU113, a residue highly conserved among vertebrates, plays a critical role in stabilizing the PSB^+^,^6^ as observed in bovine rhodopsin.^40^ In addition, GLU113 is essential for effective G protein activation, ^41^ highlighting its dual significance in opsin function. In general, counter-negative ion groups play a significant role in the electrostatic energy change upon chromophore excitation, making them key factors in determining the absorption maximum.^31,42^ Here, we found that the distances between the oxygen atoms of the GLU113 counterion and the N+ are more frequently within the ideal range for hydrogen bond formation in Rh2-1 compared to Rh2-4 (Figure S8 in the Supplementary Material). This aligns with a study suggesting that the mutual distance between the PSB and its counterion group, even with a small difference of less than 1 Å, is a key factor in controlling the absorption maximum and can lead to significant changes in the observed spectrum. ^27^ Additionally, graph-based analysis of HBs during the entire simulation showed that hydrogen bonding between E113 and the SBH+ had higher occupancy in Rh2-1 than in Rh2-4 (Figures S9 and S10 in the Supplementary Material). According to the 2D interaction diagram, the environment near SBP+ is more negatively charged in Rh2-1 than in Rh2-4. This increased negative environment could over-stabilize the positive charge on SBP+. In summary, our results demonstrate differential stability of the E113-SBP interaction between Rh2-1 and Rh2-4, with the interaction being more stable in the Rh2-1 system.

Residue 122 is known to be critical in a spectral shift of Rh1 and Rh2 from several species.^14^ Homology models of zebrafish Rh2 cone opsins in this work confirm the proximity of GLU122 to the *β*-ionone ring of the chromophore, similar to bovine rhodopsin^9^ and zebrafish rhodopsin.^19^ The key role of this site aligns with studies on zebrafish Rh2 opsins, where spectral diversification is believed to be driven by direct interactions with retinal, particularly involving residue 122 in the retinal-binding region.^20^ As has been proposed previously^43^ in 14 teleost Rh2 opsins, the negative charge on GLU122 may encourage the hydrophobic *β*-ionone ring to explore other space in the binding pocket and increase its fluctuation, resulting in a red-shift Rh2-4, whereas GLN122 in Rh2-1 discourages such fluctuations. Furthermore, here we observed that GLU122 in Rh2-4 distorts the alpha helix turn in helix 4, which leads to changes in the hydrogen bond networks around the chromophore (Figures S9 and S10 from Supplemental Material). This is similar to what was observed in a study that demonstrated that GLU122 is part of a hydrogen bond network that stabilizes the photoreceptor molecule in rhodopsin,^44^ highlighting its potential significance in Rh2 zebrafish opsins.

Lastly, site 181 has been identified as a critical factor in opsin function. The glutamic acid at position 181 is highly conserved across both vertebrate and invertebrate rhodopsins, ^36,45^ originally functioning as the sole counterion neutralizing the positive charge of the Schiff base. This ancestral role continues in invertebrates, but in vertebrates, this residue remains connected via a water-mediated hydrogen bond network to the PSB.^46^ In bovine rhodopsin, GLU181 is positioned in the extracellular loop between helices 4 and 5, oriented toward the center of the retinal’s polyene chain.^35^ A different set of studies has suggested that the spectral tuning mechanism involving GLU181 in rhodopsin is a perturbation of the electron distribution near the center of the polyene chain. ^5,16^ While GLU181 is proposed to serve as a secondary counterion during Rhodopsin activation,^34^ some studies argue that it affects spectral absorption rather than acting as a counterion. For instance, replacing GLU181 with GLN in bovine rhodopsin caused a 10 nm red-shift in the absorption maximum^36^ or has been shown to modulate the spectral properties and the stability of metarhodopsin II.^35^ However, mutational experiments indicate only minimal spectral shifts associated with this residue,^35^ and some research suggests that GLU181 primarily prevents solvent access to the Schiff base in the dark.^35^ Interestingly, GLU181 occupies different positions in the binding pockets of Rh2-1 and Rh2-4, with its placement in Rh2-4 near a water trafficking pathway not observed in Rh2-1. These differential hydration patterns lead to changes in the electrostatic environment, side-chain rotations, hydrogen bond networks, and competitive interactions between residues and water molecules. Such changes in hydration and binding pocket architecture, particularly around the PSB group, likely contribute to the distinct behaviors of these opsins. It’s also possible that residues influencing spectral shifts do so indirectly, by altering binding site conformation or the orientation of other residues. The role of water molecules in spectral tuning is further supported by structural models in rhodopsin, which suggest that conserved water networks around the retinal pocket play a regulatory role. ^27,46,47^ We propose that GLU181 is a key tuning site, influencing water molecule trafficking within the retinal-binding pocket and thereby altering its hydrophobicity or hydrophilicity, potentially affecting the protonation or deprotonation status of the Schiff base.

In quantum theory, the positive charge in the ground state of the chromophore is typically localized on the Schiff base nitrogen, while in the excited state, this charge becomes more delocalized, extending through the conjugated system towards the ionone ring. This shift in charge distribution results in an increase in the energy required for photon absorption, leading to a blue shift in the absorption spectrum.^48^ It has been suggested that blue shifts in opsins occur when polar or charged residues near the PSB preferentially stabilize the ground state over the excited state of the chromophore.^6^ Conversely, red shift models propose that polar residues along the chromophore backbone, particularly near the ionone ring, preferentially stabilize the excited state over the ground state.^49^ In our case, residue GLU122 increases chromophore motion in green-shifted pigments in the dark state, possibly raising the energy of the ground state, which in turn decreases the energy gap between the ground and excited states. This reduction in the energy difference may partly explain the higher *λ_max_* observed in our study as previously proposed in a broad set of Rh2 teleost. ^43^ Additionally, the negatively charged GLU122 in Rh2-4 tends to stay further from the negatively charged GLU181, resulting in a more distant placement of GLU181 from the chromophore. In contrast, the polar GLN122 in Rh2-1 remains closer to the ionone ring and exerts less influence on GLU181, allowing GLU181 to position itself closer to the PSB. This proximity potentially further stabilizes the ground state over the excited state in Rh2-1, leading to a blue shift.

Through QM/MM optimizations followed by excited-state calculations, we identified the electronic effects contributing to the bathochromic shift in Rh2-4 compared to Rh2-1, while also benchmarking our methodology for analyzing this system. When performing QM/MM, the importance of the QM region selection cannot be underestimated. While some studies in the literature included only chromophore^50^ terminating with the link-hydrogen-atom in place of the chromophore-bound LYS296 sidechain, or whole LYR296 residue,^51^ others consider GLU113 and water molecules in close proximity or even larger systems, involving LYR, GLU113, GLU181, TYR268, and THR94. ^52^

Our initial QM/MM optimization (Model 1), which included only chromophore and GLU113 residue in the QM region, indicated that Rh2-1 formed a unique *π*-cation interaction between the NH^+^ of the SBH^+^ and PHE293. This interaction was not present in Rh2-4. This could suggest a potential stabilizing role of PHE293 on Rh2-1’s electronic configuration, affecting its absorption properties. However, upon re-optimization (Model 2) followed by TDDFT calculations, the absorption maxima did not align with experimental trends, falsely predicting a bathochromic shift for Rh2-1. This underscored the importance of careful QM region selection, as the MM treatment of PHE293 with the less accurate LACVP* basis set may have caused incorrect optimization. Being captured in the local minimum, even after the re-optimization as a part of a QM region using the more accurate cc-pVTZ(-f) basis set, PHE293 did not change its positioning. Model 3 expanded the QM region to include additional residues (GLU113, GLU122, PHE293, GLU181, and VAL/LEU300) to address this. QM/MM optimization was carried out on the original structure with a cc-pVTZ(-f) basis set and the new QM region. This refined approach removed the artifact of the *π*-cation interaction in Rh2-1, providing a more accurate representation.

Another important observation was made in the frames of the benchmark TDDFT calculations when tested on Model 2. The calculations across various functionals and basis sets revealed that UV-VIS spectra appeared similar for all approaches except the PBE functional, which showed insufficient results for Rh2 proteins. The hybrid exchange-correlation functional CAM-B3LYP and a range-separated version of Becke’s 97 functional wB97XD provided similar and reliable results, with CAM-B3LYP being less time-consuming. The custom-developed 6-31G_JSKE basis set matched the accuracy of the significantly more time-intensive and widely used correlation-consistent polarized valence triple-zeta basis set, Dunning’s cc-pVTZ, establishing it as a viable choice. Interestingly, gas-phase calculations displayed a larger *λ*_max_ difference (36-37 nm) compared to those in water solvent (21-23 nm), with the exception of HF/3-21G, which produced a similar Δ*λ*_max_ as the water phase calculations (*≈* 25 nm). This indicated that solvent effects play a significant role in accurately modeling opsin-chromophore systems, justifying CAM-B3LYP/6-31G_JSKE in a water solvent for subsequent studies.

The HOMO-LUMO gap is often correlated with the absorption maximum. This gap represents the energy difference between the ground state (HOMO) and the excited state (LUMO), which corresponds to the energy required for an electronic transition when the molecule absorbs light. This correlation was reaffirmed, as Rh2-1’s larger gap reflected shorter wavelengths, while Rh2-4 exhibited a smaller gap and a bathochromic shift. Notably, Rh2-4 maintained consistent HOMO and LUMO positioning across all methods, while Rh2-1 displayed variability. The HF/3-21G level of theory positioned HOMO solely on GLU181, yielding a more negative HOMO energy and, thus, a larger HOMO-LUMO gap. In this case, on the UV-Vis spectrum, the more significant gap between *λ*_max_ was noted (15 nm), which correlated better with the experimental data (Δ*λ*_max_ = 38 nm). Contrary to this, both QM/MM optimization with B3LYP/cc-pVTZ(-f) and QM TDDFT calculations with CAM-B3LYP/6-31G_JSKE showed a delocalized HOMO between the chromophore and GLU181, resulting in a higher energy HOMO and, thus, reduced gap. Additional calculations were done using CAM-B3LYP/cc-pVTZ to verify this, which confirmed the latest results. The Hartree-Fock method neglects electron correlation effects, crucial for accurately describing electronic interactions, especially in conjugated and complex systems such as Rh2. In addition to that, the lack of polarization and diffuse functions in 3-21G makes it challenging to accurately represent delocalized electrons, especially in interactions involving charged species. Finally, the gas-phase calculation does not account for the solvent environment, which can significantly influence the electronic structure when charged and polar species are present. Thus, while HF/3-21G successfully predicted the direction of the absorption maximum shift and provided a more apparent distinction between the two pigments, it did not perform well with either the size of this shift or the electronic structure of the chromophore. The noteworthy discrepancies in the HOMO-LUMO gap between methods (8.24 eV vs. 5.37 eV) and the difference in absorption maxima highlight the impact of computational choices on electronic transition interpretations.

A critical challenge was to address why the absorption maximum shift was significantly smaller than the experimental results. TDDFT predictions often require scaling to align with experimental data.^53^ Considering this, it was challenging to conclude whether the calculated value for Rh2-1 was extensively red-shifted or the one for Rh2-4 was far blue-shifted. The answer to this question could be speculated based on the location of HOMO of Rh2-4 being located on a GLU122. As was shown earlier, the distance between the chromophore and GLU122 fluctuated significantly throughout the MD simulation time. With such fluctuations, when the distance between this residue and LYR increases, it destabilizes the ground state, resulting in higher energy of HOMO. The energy of LUMO, on the contrary, does not depend on this distance; thus, even considering the flexibility of a chromophore, it remains mostly unchanged. The persistent increase in HOMO energy while keeping LUMO energy relatively stable could result in a smaller average HOMO-LUMO gap and a bathochromic shift consistent with experimental observations.

Biophysically relevant questions often focus on subtle molecular differences between closely related states or systems. This can include identifying molecular signatures when a protein binds to different ligands, evaluating the effects of point mutations on the same protein, or, as in this case, analyzing the same chromophore bound in different proteins. Traditionally, these analyses involve the manual interpretation of large, high-dimensional data sets, requiring both expertise and a keen eye that has become almost an art form. Therefore, the complexity of such tasks underscores the growing need for automated methods to streamline the selection and prioritization of key features for more efficient analysis. Future efforts should focus on applying machine learning algorithms to advance this process further.

## Conclusions

This study employed molecular modeling techniques to investigate the mechanisms behind the spectral shifts in zebrafish Rh2-1 and Rh2-4 cone opsins. Through all-atom molecular dynamics simulations and structural analyses, we identified key amino acid sites that contribute to the distinct spectral properties of these pigments. The comparison between Rh2-1 and Rh2-4 revealed differential stability in key interactions, particularly between GLU113 GLN/GLU122 and GLU181 and the protonated Schiff base, which we propose as drivers of the observed spectral shift. Further investigation into the hydrogen bonding and electrostatic environments around critical residues highlights their potential role in tuning absorption maxima. The QM calculations revealed that fluctuations in chromophore-GLU122 distance could be one of the driving forces of the bathochromic shift in Rh2-4, as this pigment’s highest occupied molecular orbital was located on GLU122. Our findings provide valuable insights into the chromophore-protein interactions that govern spectral tuning in visual pigments, suggesting that these residues and water-mediated interactions play a critical role in the photoreceptor function of zebrafish opsins.

## Methods

The amino acid sequences of the zebrafish (Danio rerio) Rh2-1 and Rh2-4 cone opsins^13,20^ were obtained from UniProt with accession numbers Q9W6A5 and Q9W6A6, respectively. A template structure search was performed using MODELLER v9.15.^54^ The bovine rhodopsin (Rh1) structure (PDB ID: 1U19)^10^ was selected as the template for both opsins because it met the following criteria: i) sequence identity >60% (Table 1); ii) >95% sequence coverage with the target Rh2 sequence; iii) presence of 11-*cis* retinal bound in the binding pocket and occupied palmitoylation sites; iv) high X-ray crystal resolution (2.2 Å); and v) no mutations in the crystal structure protein. MODELLER v9.15 was used to perform the sequence alignments and generate three-dimensional structures of the Rh2 cone opsins. Five homology structures were generated for each opsin sequence. Stereochemical checks were performed on all five structures using the SWISS-MODEL structure evaluation tool (https://swissmodel.expasy.org/), and the best structure was selected based on minimal stereochemical deviation and a high QMEAN score.

The final selected structure of each Rh2 opsin was first uploaded to the Prediction of Proteins in Membranes web server (http://opm.phar.umich.edu/server.php). The membrane boundaries provided by this server, along with the protein model, were then uploaded to the CHARMM-GUI server (http://charmm-gui.org/) for further processing. To incorporate the 11-*cis* retinal chromophore into the binding pocket of each structure, the three-letter amino acid code of the lysine residue that covalently binds to the chromophore to form the Schiff base was changed from LYS to LYR in the PDB file. This modification allowed CHARMM-GUI to recognize and build the Schiff base using the appropriate force field parameters available on the server. The palmitate moiety was included because the sequences share a conserved palmitoylation site (C323) with the bovine rhodopsin template sequence. The C323 residue was bound to a palmitate molecule using the “add palmitoylation sites” option in CHARMM-GUI. Protonation states of the amino acid residues were assigned at the physiological pH of 7.4. The protein was embedded in an unsaturated homogeneous bilayer of 1-stearoyl-2-docosahexaenoyl-sn-glycero-3-phosphocholine (SDPC) lipids to provide a realistic representation of the phospholipids found in the cone outer segment. The replacement method^55^ was used to pack the opsin model with the lipid bilayer, with a lipid layer thickness of 1.6 ( 70 lipids in the top leaflet and 70 lipids in the bottom leaflet). Each system was placed in a rectangular solvent box, and a 10 Å TIP3P water layer was added to solvate the intra– and extracellular spaces. Charge neutrality was achieved by adding Na^+^ and Cl*^−^* ions at a concentration of 0.15 mol/L to the water layers. CHARMM-GUI initially assumed the retinal was in the 11-*trans* conformation, so we replaced the coordinates with the 11-*cis* conformation obtained from the bovine rhodopsin structure. The CHARMM36 force field^56^ parameters were used for the protein and lipids. Each system was first minimized using the steepest descent for 5,000 steps. To allow equilibration of the water, each system was then simulated for a total of 550 ps with harmonic restraints on all heavy atoms in the protein, phosphorus atoms in the lipid head group, and all dihedral angles in the lipid carbon chains. The restrained simulations were divided into six steps, with restraints gradually relaxed at each step. The temperature was set to 300 K, and the pressure was maintained at 1 atm using the Berendsen algorithm. Production NPT simulations were carried out for 100 ns using the Parrinello-Rahman barostat^57^ with semi-isotropic pressure coupling and the Nose-Hoover thermostat^58^ for temperature control.

The analysis was performed within Python Jupyter notebooks. ^59^ MDAnalysis packages were used for the majority of structural feature calculations on the MD trajectories.^60^ Sequence alignment was performed using MultAlin software. ^61^ Bridge2 software was employed for HBs graph-based network analysis. ^62^ Maestro software from Schrödinger was used for 2D interaction diagrams. ^21^

The structures of Rh2-1 and Rh2-4 were optimized for QM/MM calculations using the Qsite module,^63^ implemented in the Schrödinger Software Package. The MM region was minimized using the Truncated Newton algorithm with the OPLS_2005 force field. The importance of carefully selecting the QM region in QM/MM studies cannot be overstated, as it significantly affects the accuracy of electronic properties. We established three QM/MM models: (1) the preliminary model 1 utilized the B3LYP functional^64^ with the LACVP* basis set,^65^ incorporating the entire GLU113 and LYS296 residues in the QM region, while the remainder of the protein was treated as the MM region; (2) the initial geometry from model 1 refined using the B3LYP functional with the cc-pVTZ(-f) basis set^66^ and including in the QM region the entire LYR296 residue and the side chains of GLU113 and PHE293; and (3) the original geometry optimized using B3LYP/cc-pVTZ(-f) with LYR296 as the entire residue, and GLU181, GLU113, GLN(GLU)122, PHE293, and VAL(LEU)300 as side chains in the QM region. HOMO and LUMO orbitals were calculated for these optimized structures. QM calculations were carried out for optimized structures from models 2 and 3 using Gaussian 09, utilizing single-point calculations and Time-Dependent Density Functional Theory (TDDFT).^67–69^ For model 2, we selected all the residues that interacted with the chromophore and those that interacted with the above-mentioned residues. This way, the structure for QM calculations contained LYR296, TYR43, SER(THR)94, GLU113, ALA117, SER186, SER187, PHE293, THR297, and VAL300 (213 atoms for Rh2-1 and 219 atoms for Rh2-4). Various computational methods were used, including Hartree-Fock (HF),^70^ PBE,^71^ wB97XD,^72^ and CAM-B3LYP.^73^ A range of basis sets was employed, including 3-21G,^74^ 6-31G_JSKE,^22^ and cc-pVTZ.^66^ Calculations were carried out in both the gas phase and the conductor-like polarizable continuum model (CPCM)^75,76^ with water as the solvent. This allowed us to perform a benchmark, which suggested using only HF/3-21G in the gas phase and CAM-B3LYP/6-31G_JSKE in the water solvent for future calculations. For Model 3, the QM calculations involved LYR296, GLU181, GLU113, GLN(GLU)122, PHE293, and VAL(LEU)300 (163 atoms for Rh2-1 and 164 atoms for Rh2-4). Cubes were generated using a standalone Cubegen program from the Gaussian 09 package.

## Supporting information

Supporting Information Available

## Acknowledgement

LAC, SKP, PLA, JEB and JSP were supported by the COBRE Research Project Grant to JSP from National Institute of General Medical Sciences (NIGMS) of the National Institutes of Health (NIH) under Award Number P20GM104420. Computational resources were provided in part by Research Computing and Data Services in the Institute for Interdisciplinary Data Science at University of Idaho. K.K was partially supported by the Mississippi IN-BRE, funded by an Institutional Development Award (IDeA) from the NIGMS of the NIH under grant number P20GM103476 and by the National Science Foundation award number OIA-2414445. The content is solely the responsibility of the authors and does not necessarily represent the official views of the funding agencies.

## Notes

### Competing Interest Statement

The authors have declared no competing interest.

## References

(1) Karasuyama, M., Inoue, K., Nakamura, R., Kandori, H., and Takeuchi, I. (2018) Understanding colour tuning rules and predicting absorption wavelengths of microbial rhodopsins by data-driven machine-learning approach. Scientific reports 8, 15580.

(2) Shichida, Y., and Matsuyama, T. (2009) Evolution of opsins and phototransduction. Philosophical Transactions of the Royal Society B: Biological Sciences 364, 2881–2895.

(3) Nakanishi, K. (1991) Why 11-cis-retinal? American Zoologist 31, 479–489.

(4) Honig, B., Greenberg, A. D., Dinur, U., and Ebrey, T. G. (1976) Visual-pigment spectra: implications of the protonation of the retinal Schiff base. Biochemistry 15, 4593–4599.

(5) Honig, B., Dinur, U., Nakanishi, K., Balogh-Nair, V., Gawinowicz, M. A., Arnaboldi, M., and Motto, M. G. (1979) An external point-charge model for wavelength regulation in visual pigments. Journal of the American Chemical Society 101, 7084– 7086.

(6) Chang, B. S., Crandall, K. A., Carulli, J. P., and Hartl, D. L. (1995) Opsin phylogeny and evolution: a model for blue shifts in wavelength regulation. Molecular phylogenetics and evolution 4, 31–43.

(7) Lin, S. W., Kochendoerfer, G. G., Carroll, K. S., Wang, D., Mathies, R. A., and Sakmar, T. P. (1998) Mechanisms of spectral tuning in blue cone visual pigments: visible and raman spectroscopy of blue-shifted rhodopsin mutants. Journal of Biological Chemistry 273, 24583–24591.

(8) Lagman, D., Ocampo Daza, D., Widmark, J., Abalo, X. M., Sundström, G., and Larhammar, D. (2013) The vertebrate ancestral repertoire of visual opsins, transducin alpha subunits and oxytocin/vasopressin receptors was established by duplication of their shared genomic region in the two rounds of early vertebrate genome duplications. BMC evolutionary biology 13, 1–21.

(9) Palczewski, K., Kumasaka, T., Hori, T., Behnke, C. A., Motoshima, H., Fox, B. A., Trong, I. L., Teller, D. C., Okada, T., Stenkamp, R. E., and others (2000) Crystal structure of rhodopsin: AG protein-coupled receptor. science 289, 739–745.

(10) Okada, T., Sugihara, M., Bondar, A.-N., Elstner, M., Entel, P., and Buss, V. (2004) The retinal conformation and its environment in rhodopsin in light of a new 2.2 Å crystal structure. Journal of molecular biology 342, 571–583.

(11) Peng, Q., Li, J., Jiang, H., Cheng, X., Lu, Q., Zhou, S., Zhang, Y., Lv, S., Wan, S., Yang, T., and others (2024) Cryo-EM structures of human cone visual pigments. bioRxiv 2024–01.

(12) Fadool, J. M., and Dowling, J. E. (2008) Zebrafish: a model system for the study of eye genetics. Progress in retinal and eye research 27, 89–110.

(13) Chinen, A., Hamaoka, T., Yamada, Y., and Kawamura, S. (2003) Gene duplication and spectral diversification of cone visual pigments of zebrafish. Genetics 163, 663–675.

(14) Takahashi, Y., and Ebrey, T. G. (2003) Molecular basis of spectral tuning in the newt short wavelength sensitive visual pigment. Biochemistry 42, 6025–6034.

(15) Imai, H., Kojima, D., Oura, T., Tachibanaki, S., Terakita, A., and Shichida, Y. (1997) Single amino acid residue as a functional determinant of rod and cone visual pigments. Proceedings of the National Academy of Sciences 94, 2322–2326.

(16) Menon, S. T., Han, M., and Sakmar, T. P. (2001) Rhodopsin: structural basis of molecular physiology. Physiological reviews 81, 1659–1688.

(17) Pogozheva, I. D., Lomize, A. L., and Mosberg, H. I. (1997) The transmembrane 7-alpha-bundle of rhodopsin: distance geometry calculations with hydrogen bonding constraints. Biophysical Journal 72, 1963–1985.

(18) Yokoyama, S., Zhang, H., Radlwimmer, F. B., and Blow, N. S. (1999) Adaptive evolution of color vision of the Comoran coelacanth (Latimeria chalumnae). Proceedings of the National Academy of Sciences 96, 6279–6284.

(19) Morrow, J. M. Molecular evolution of zebrafish (Danio rerio) visual pigment function; University of Toronto (Canada), 2014.

(20) Chinen, A., Matsumoto, Y., and Kawamura, S. (2005) Reconstitution of ancestral green visual pigments of zebrafish and molecular mechanism of their spectral differentiation. Molecular biology and evolution 22, 1001–1010.

(21) Maestro, S. (2020) Schrödinger Release 2015-1: Maestro. Schrödinger, LLC, New York, NY 2020.

(22) Kapusta, K., Sizochenko, N., Karabulut, S., Okovytyy, S., Voronkov, E., and Leszczynski, J. (2018) QSPR modeling of optical rotation of amino acids using specific quantum chemical descriptors. Journal of Molecular Modeling 24.

(23) Kong, Y., and Karplus, M. (2007) The signaling pathway of rhodopsin. Structure 15, 611–623.

(24) Zabelskii, D., Dmitrieva, N., Volkov, O., Shevchenko, V., Kovalev, K., Balandin, T., Soloviov, D., Astashkin, R., Zinovev, E., Alekseev, A., and others (2021) Structurebased insights into evolution of rhodopsins. Communications biology 4, 821.

(25) Fritze, O., Filipek, S., Kuksa, V., Palczewski, K., Hofmann, K. P., and Ernst, O. P. (2003) Role of the conserved NPxxY (x) 5, 6F motif in the rhodopsin ground state and during activation. Proceedings of the National Academy of Sciences 100, 2290–2295.

(26) Zhou, X. E., Melcher, K., and Xu, H. E. (2012) Structure and activation of rhodopsin. Acta Pharmacologica Sinica 33, 291–299.

(27) Fujimoto, K., Hayashi, S., Hasegawa, J.-y., and Nakatsuji, H. (2007) Theoretical studies on the color-tuning mechanism in retinal proteins. Journal of chemical theory and computation 3, 605–618.

(28) Wanko, M., Hoffmann, M., Strodel, P., Koslowski, A., Thiel, W., Neese, F., Frauenheim, T., and Elstner, M. (2005) Calculating absorption shifts for retinal proteins: computational challenges. The Journal of Physical Chemistry B 109, 3606–3615.

(29) Andruniów, T., Ferré, N., and Olivucci, M. (2004) Structure, initial excited-state relaxation, and energy storage of rhodopsin resolved at the multiconfigurational perturbation theory level. Proceedings of the National Academy of Sciences 101, 17908–17913.

(30) Sugihara, M., Hufen, J., and Buss, V. (2006) Origin and consequences of steric strain in the rhodopsin binding pocket. Biochemistry 45, 801–810.

(31) Hayashi, S., Tajkhorshid, E., Pebay-Peyroula, E., Royant, A., Landau, E. M., Navarro, J., and Schulten, K. (2001) Structural determinants of spectral tuning in retinal proteins bacteriorhodopsin vs sensory rhodopsin II. The Journal of Physical Chemistry B 105, 10124–10131.

(32) Fujimoto, K., Hasegawa, J.-y., Hayashi, S., Kato, S., and Nakatsuji, H. (2005) Mechanism of color tuning in retinal protein: SAC-CI and QM/MM study. Chemical physics letters 414, 239–242.

(33) Cohen, G. B., Oprian, D. D., and Robinson, P. R. (1992) Mechanism of activation and inactivation of opsin: role of Glu113 and Lys296. Biochemistry 31, 12592–12601.

(34) Lüdeke, S., Beck, M., Yan, E. C., Sakmar, T. P., Siebert, F., and Vogel, R. (2005) The role of Glu181 in the photoactivation of rhodopsin. Journal of molecular biology 353, 345–356.

(35) Yan, E. C., Kazmi, M. A., De, S., Chang, B. S., Seibert, C., Marin, E. P., Mathies, R. A., and Sakmar, T. P. (2002) Function of extracellular loop 2 in rhodopsin: glutamic acid 181 modulates stability and absorption wavelength of metarhodopsin II. Biochemistry 41, 3620–3627.

(36) Terakita, A., Yamashita, T., and Shichida, Y. (2000) Highly conserved glutamic acid in the extracellular IV–V loop in rhodopsins acts as the counterion in retinochrome, a member of the rhodopsin family. Proceedings of the National Academy of Sciences 97, 14263–14267.

(37) Blatz, P. E., Mohler, J. H., and Navangul, H. V. (1972) Anion-induced wavelength regulation of absorption maxima of Schiff bases of retinal. Biochemistry 11, 848–855.

(38) Kakitani, H., Kakitani, T., Rodman, H., and Honig, B. (1985) On the mechanism of wavelength regulation in visual pigments. Photochemistry and photobiology 41, 471–479.

(39) Irving, C. S., Byers, G. W., and Leermakers, P. A. (1970) Spectroscopic model for the visual pigments. Influence of microenvironmental polarizability. Biochemistry 9, 858–864.

(40) Sakmar, T. P., Franke, R. R., and Khorana, H. G. (1989) Glutamic acid-113 serves as the retinylidene Schiff base counterion in bovine rhodopsin. Proceedings of the National Academy of Sciences 86, 8309–8313.

(41) Terakita, A., Koyanagi, M., Tsukamoto, H., Yamashita, T., Miyata, T., and Shichida, Y. (2004) Counterion displacement in the molecular evolution of the rhodopsin family. Nature structural & molecular biology 11, 284–289.

(42) Schreiber, M., Buß, V., and Sugihara, M. (2003) Exploring the Opsin shift with ab initio methods: Geometry and counterion effects on the electronic spectrum of retinal. The Journal of chemical physics 119, 12045–12048.

(43) Patel, J. S., Brown, C. J., Ytreberg, F. M., and Stenkamp, D. L. (2018) Predicting peak spectral sensitivities of vertebrate cone visual pigments using atomistic molecular simulations. PLOS Computational Biology 14, e1005974.

(44) Patel, A. B., Crocker, E., Reeves, P. J., Getmanova, E. V., Eilers, M., Khorana, H. G., and Smith, S. O. (2005) Changes in interhelical hydrogen bonding upon rhodopsin activation. Journal of molecular biology 347, 803–812.

(45) Porter, M. L., Blasic, J. R., Bok, M. J., Cameron, E. G., Pringle, T., Cronin, T. W., and Robinson, P. R. (2012) Shedding new light on opsin evolution. Proceedings of the Royal Society B: Biological Sciences 279, 3–14.

(46) Gruhl, T., Weinert, T., Rodrigues, M. J., Milne, C. J., Ortolani, G., Nass, K., Nango, E., Sen, S., Johnson, P. J., Cirelli, C., and others (2023) Ultrafast structural changes direct the first molecular events of vision. Nature 615, 939–944.

(47) Okada, T., Fujiyoshi, Y., Silow, M., Navarro, J., Landau, E. M., and Shichida, Y. (2002) Functional role of internal water molecules in rhodopsin revealed by X-ray crystallography. Proceedings of the National Academy of Sciences 99, 5982–5987.

(48) Mathies, R., and Stryer, L. (1976) Retinal has a highly dipolar vertically excited singlet state: implications for vision. Proceedings of the National Academy of Sciences 73, 2169–2173.

(49) Chan, T., Lee, M., and Sakmar, T. (1992) Introduction of hydroxyl-bearing amino acids causes bathochromic spectral shifts in rhodopsin. Amino acid substitutions responsible for red-green color pigment spectral tuning. Journal of Biological Chemistry 267, 9478– 9480.

(50) Orozco-Gonzalez, Y., Manathunga, M., Marín, M. d. C., Agathangelou, D., Jung, K.- H., Melaccio, F., Ferré, N., Haacke, S., Coutinho, K., Canuto, S., and Olivucci, M. (2017) An Average Solvent Electrostatic Configuration Protocol for QM/MM Free Energy Optimization: Implementation and Application to Rhodopsin Systems. Journal of Chemical Theory and Computation 13, 6391–6404.

(51) Borin, V. A., Wiebeler, C., and Schapiro, I. (2018) A QM/MM study of the initial excited state dynamics of green-absorbing proteorhodopsin. Faraday Discussions 207, 137–152.

(52) Hernández-Rodríguez, E. W., Sánchez-García, E., Crespo-Otero, R., Montero-Alejo, A. L., Montero, L. A., and Thiel, W. (2012) Understanding Rhodopsin Mutations Linked to the Retinitis pigmentosa Disease: a QM/MM and DFT/MRCI Study. The Journal of Physical Chemistry B 116, 1060–1076.

(53) Bremond, E. A., Kieffer, J., and Adamo, C. (2010) A reliable method for fitting TD-DFT transitions to experimental UV–visible spectra. Journal of Molecular Structure: THEOCHEM 954, 52–56.

(54) Webb, B., and Sali, A. (2016) Comparative protein structure modeling using MODELLER. Current protocols in bioinformatics 54, 5–6.

(55) Woolf, T. B., and Roux, B. (1994) Molecular dynamics simulation of the gramicidin channel in a phospholipid bilayer. Proceedings of the National Academy of Sciences 91, 11631–11635.

(56) Huang, J., and MacKerell Jr, A. D. (2013) CHARMM36 all-atom additive protein force field: Validation based on comparison to NMR data. Journal of computational chemistry 34, 2135–2145.

(57) Parrinello, M., and Rahman, A. (1981) Polymorphic transitions in single crystals: A new molecular dynamics method. Journal of Applied physics 52, 7182–7190.

(58) Evans, D. J., and Holian, B. L. (1985) The nose–hoover thermostat. The Journal of chemical physics 83, 4069–4074.

(59) Kluyver, T., Ragan-Kelley, B., Pérez, F., Granger, B., Bussonnier, M., Frederic, J., Kelley, K., Hamrick, J., Grout, J., Corlay, S., Ivanov, P., Avila, D., Abdalla, S., and Willing, C. Jupyter Notebooks – a publishing format for reproducible computational workflows. 2016.

(60) Michaud-Agrawal, N., Denning, E. J., Woolf, T. B., and Beckstein, O. (2011) MD Analysis: a toolkit for the analysis of molecular dynamics simulations. Journal of computational chemistry 32, 2319–2327.

(61) Corpet, F. (1988) Multiple sequence alignment with hierarchical clustering. Nucleic acids research 16, 10881–10890.

(62) Siemers, M., Lazaratos, M., Karathanou, K., Guerra, F., Brown, L. S., and Bondar, A.-N. (2019) Bridge: A graph-based algorithm to analyze dynamic H-bond networks in membrane proteins. Journal of chemical theory and computation 15, 6781–6798.

(63) Murphy, R. B., Philipp, D. M., and Friesner, R. A. (2000) A mixed quantum mechanics/molecular mechanics (QM/MM) method for large-scale modeling of chemistry in protein environments. Journal of Computational Chemistry 21, 1442–1457.

(64) Becke, A. D. (1993) Density-functional thermochemistry. III. The role of exact exchange. The Journal of Chemical Physics 98, 5648–5652.

(65) Hay, P. J., and Wadt, W. R. (1985) Ab initio effective core potentials for molecular calculations. Potentials for K to Au including the outermost core orbitals. The Journal of Chemical Physics 82, 299–310.

(66) Dunning, T. H. (1989) Gaussian basis sets for use in correlated molecular calculations. I. The atoms boron through neon and hydrogen. The Journal of Chemical Physics 90, 1007–1023.

(67) Bauernschmitt, R., and Ahlrichs, R. (1996) Treatment of electronic excitations within the adiabatic approximation of time dependent density functional theory. Chemical Physics Letters 256, 454–464.

(68) Casida, M. E., Jamorski, C., Casida, K. C., and Salahub, D. R. (1998) Molecular excitation energies to high-lying bound states from time-dependent density-functional response theory: Characterization and correction of the time-dependent local density approximation ionization threshold. The Journal of Chemical Physics 108, 4439–4449.

(69) Adamo, C., and Jacquemin, D. (2013) The calculations of excited-state properties with Time-Dependent Density Functional Theory. Chem. Soc. Rev. 42, 845–856.

(70) Pople, J. A., and Nesbet, R. K. (1954) Self-Consistent Orbitals for Radicals. The Journal of Chemical Physics 22, 571–572.

(71) Perdew, J. P., Burke, K., and Ernzerhof, M. (1996) Generalized Gradient Approximation Made Simple. Physical Review Letters 77, 3865–3868.

(72) Chai, J.-D., and Head-Gordon, M. (2008) Long-range corrected hybrid density functionals with damped atom–atom dispersion corrections. Physical Chemistry Chemical Physics 10, 6615.

(73) Yanai, T., Tew, D. P., and Handy, N. C. (2004) A new hybrid exchange–correlation functional using the Coulomb-attenuating method (CAM-B3LYP). Chemical Physics Letters 393, 51–57.

(74) Binkley, J. S., Pople, J. A., and Hehre, W. J. (1980) Self-consistent molecular orbital methods. 21. Small split-valence basis sets for first-row elements. Journal of the American Chemical Society 102, 939–947.

(75) Barone, V., and Cossi, M. (1998) Quantum Calculation of Molecular Energies and Energy Gradients in Solution by a Conductor Solvent Model. The Journal of Physical Chemistry A 102, 1995–2001.

(76) Cossi, M., Rega, N., Scalmani, G., and Barone, V. (2003) Energies, structures, and electronic properties of molecules in solution with the C-PCM solvation model. Journal of Computational Chemistry 24, 669–681.

