## Supporting Information Available for "Molecular Mechanisms Underlying the Spectral Shift in Zebrafish Cone Opsins"

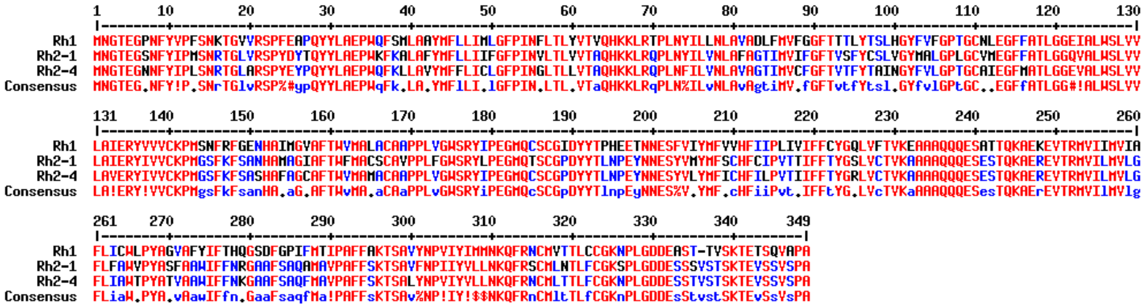

Figure S1: Sequence alignment of blue sensitive opsin (Rh2-1) and green sensitive one (Rh2-4) with bovine rhodopsin (Rh1). Rh2-1 and Rh2-4 sequences share 83% similarity; Rh1 and Rh2-4 share 71% similarity and Rh1 and Rh2-1 share 66% similarity. Alignment performed using MultAlin software.[1]

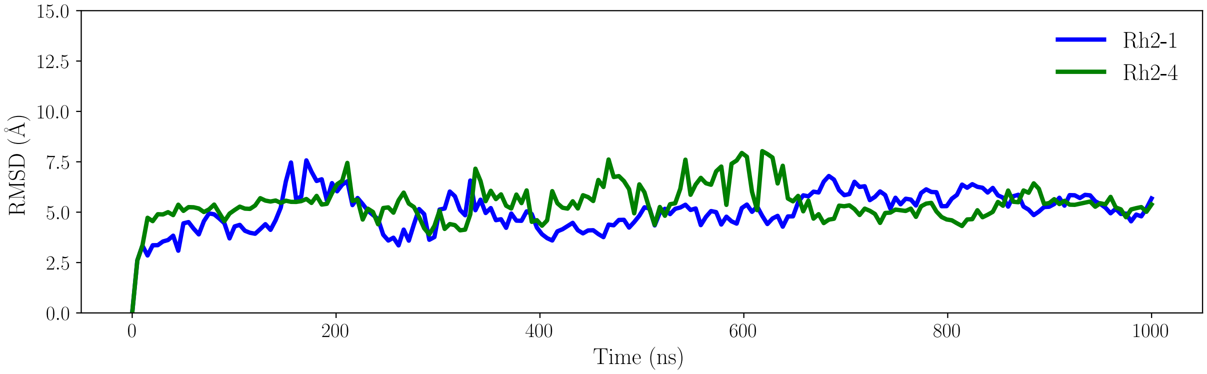


Figure S2: C$\alpha$ atoms' Root Mean Square Deviation (RMSD) for Rh2-1 and Rh2-4.

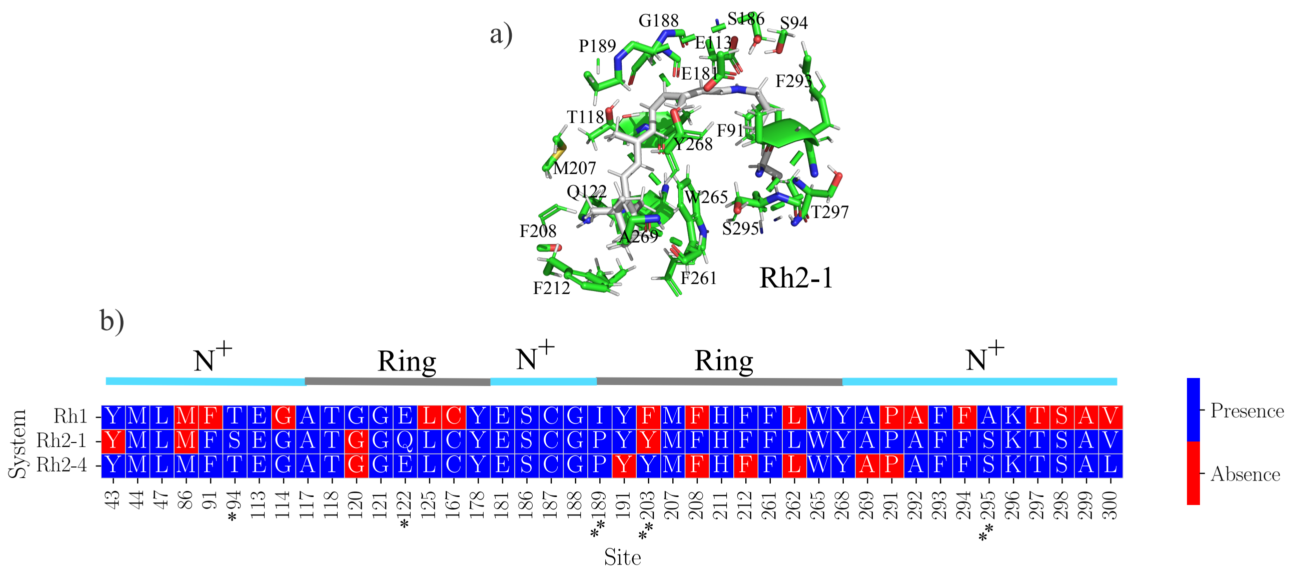


Figure S3: Residues present in the binding pocket of Rh1, Rh2-1, and Rh2-4 are detailed here. a) Residues belonging to the Rh2-1 retinal pocket in their 3D structural distribution around the chromophore. b) Comparison of sites and residues present in every system. The residues for Rh1 were sourced from reference.[2] For Rh2-1 and Rh2-4, residues were selected based on their proximity—within 4.5 Å (consistent with the threshold used for Rh1) to the chromophore in a representative structure. The presence or absence identifier specifies whether the residue at this site is included in the binding pocket for each respective system. Asterisks indicate deviations from the Rh1 sequence: one asterisk denotes that at least one system, either Rh2-1 or Rh2-4, features a different residue at that position, while two asterisks indicate that both systems have a different residue at the corresponding site. The top blue and grey lines in the chart indicate spatial proximity to either the protonated nitrogen ($N^+$) or the ring, with blue lines representing closer proximity to the $N^+$ and grey lines indicating closer proximity to the ring.

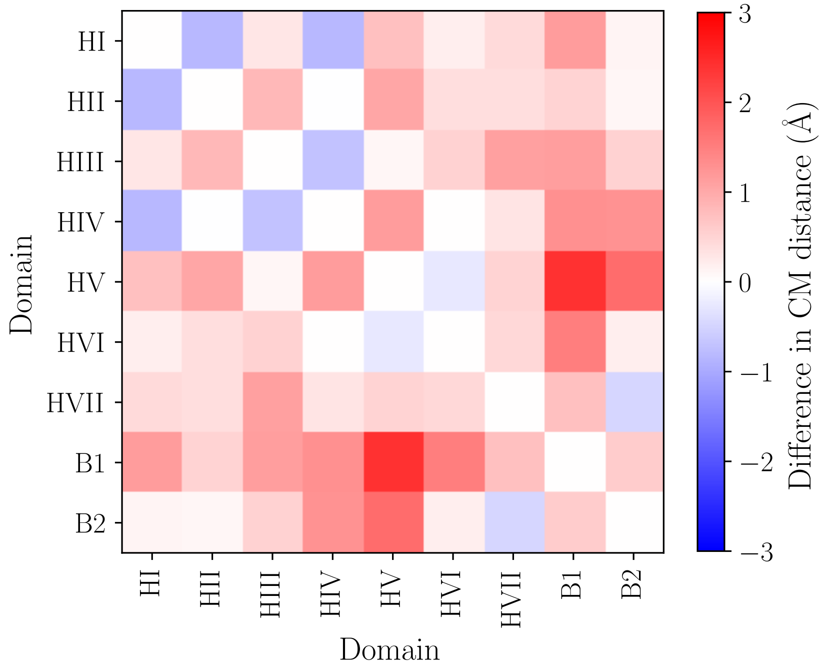


Figure S4: Distance between centers of mass of different domains on Rh2-4 (green-light sensitive) and Rh2-1 (blue-light sensitive) pigments. Blue indicates that the average distance between the centers of mass of these domains is closer in Rh2-4 compared to Rh2-1, while red indicates that they are farther apart.

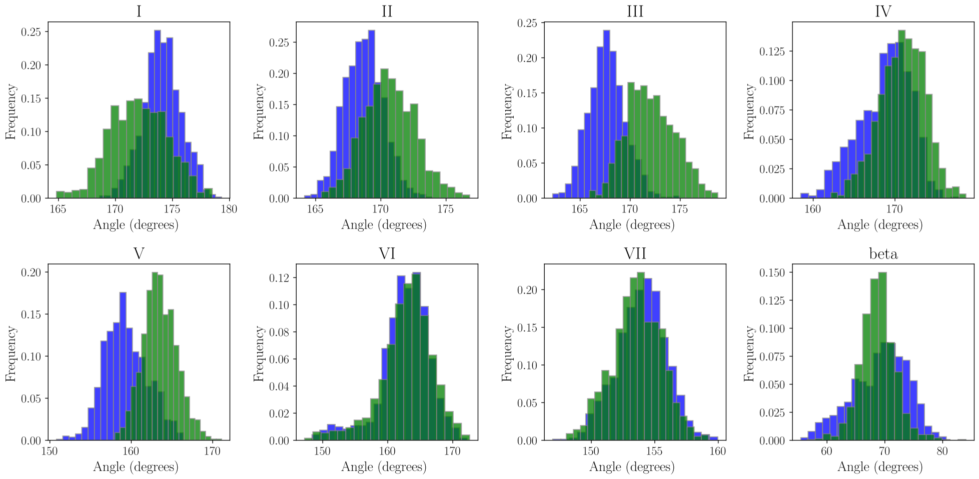


Figure S5: Geometric angle (First residue - center of mass - last residue) of each helices in Rh2-4 (green) and Rh2-1 (blue) pigments.

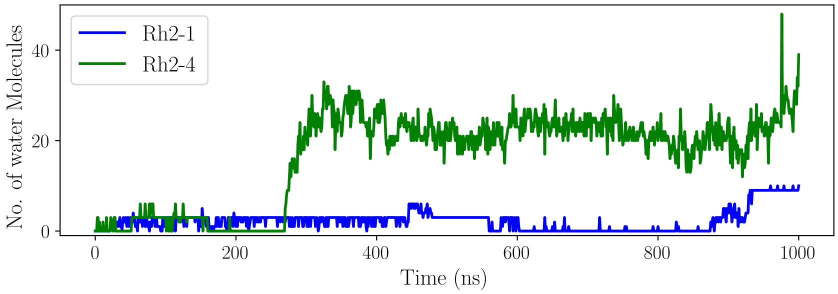


Figure S6: Hydration of the LYR pocket.

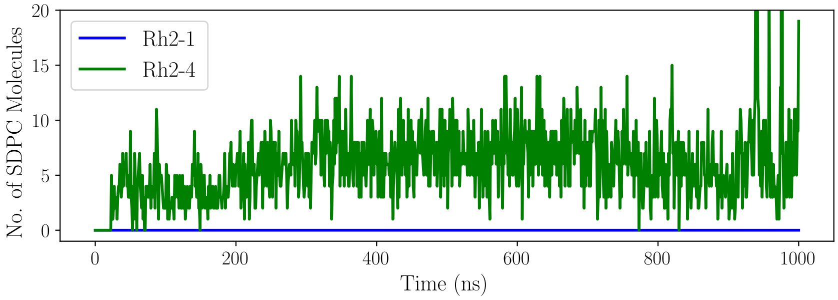


Figure S7: SDPC near the chromophore.

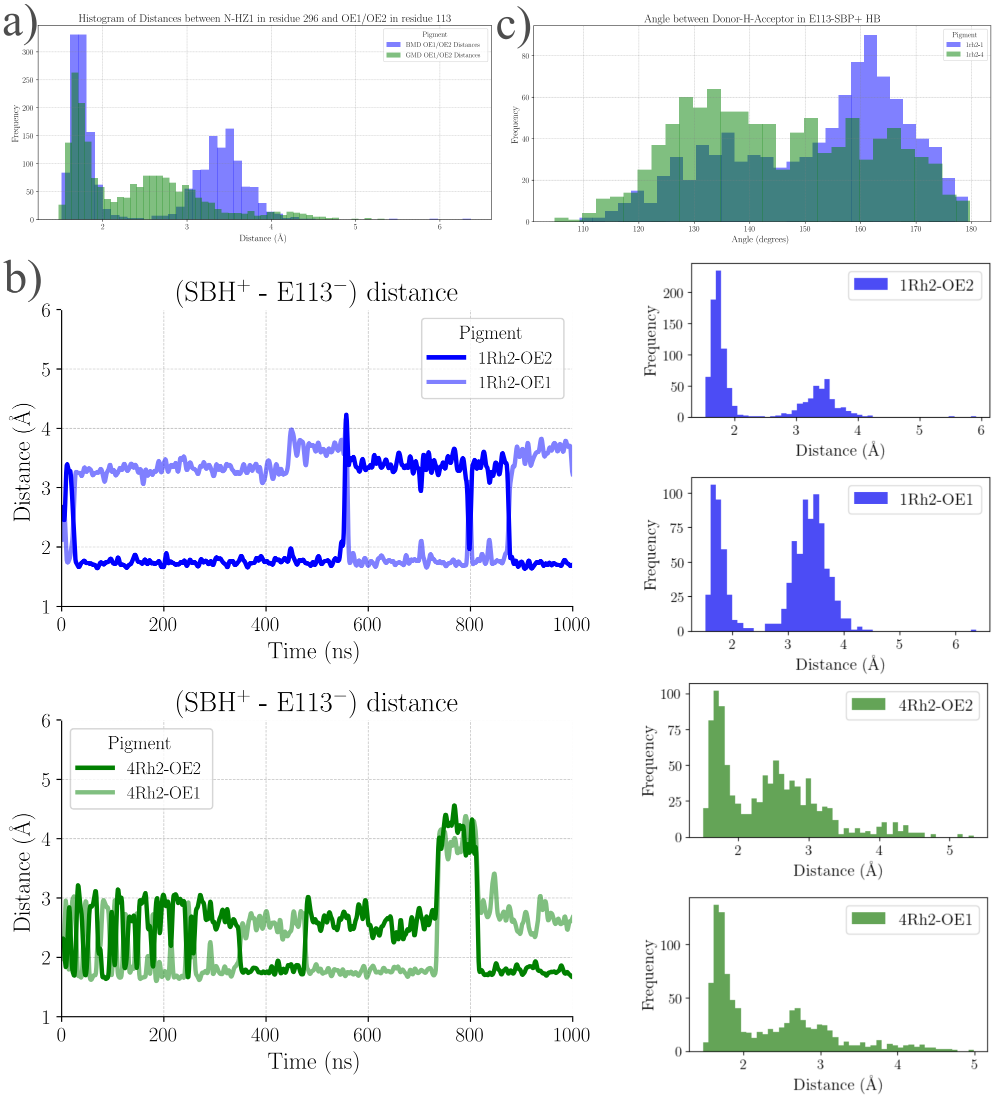


Figure S8: Analysis of GLU113-SBP$^+$ distances and donor-Hacceptor angles in Rh2-1 (1Rh2) and Rh2-4 (4Rh2). a) Distance between the SBH+ and the E113 counterion (OE2 or OE1) over time for the 1Rh2 and 4Rh2 pigments. b) Distance between the Schiff Base proton (SBH$^+$) and the E113- counterion over time for the 1Rh2 and 4Rh2 pigments. The 1Rh2 pigment shows a more stable and consistently shorter distance, suggesting a stronger or more stable interaction. In regions where the distance exceeds 3.5 Å, there is likely a water molecule interacting with the carboxylate group of E113. The two main states in the green case depend on whether the carbonyl or carboxylate oxygen interacts with the SBH$^+$. c) Angles between the donor-H-acceptor in GLU113-SBP$^+$ HB over time for the 1Rh2 and 4Rh2 pigments.

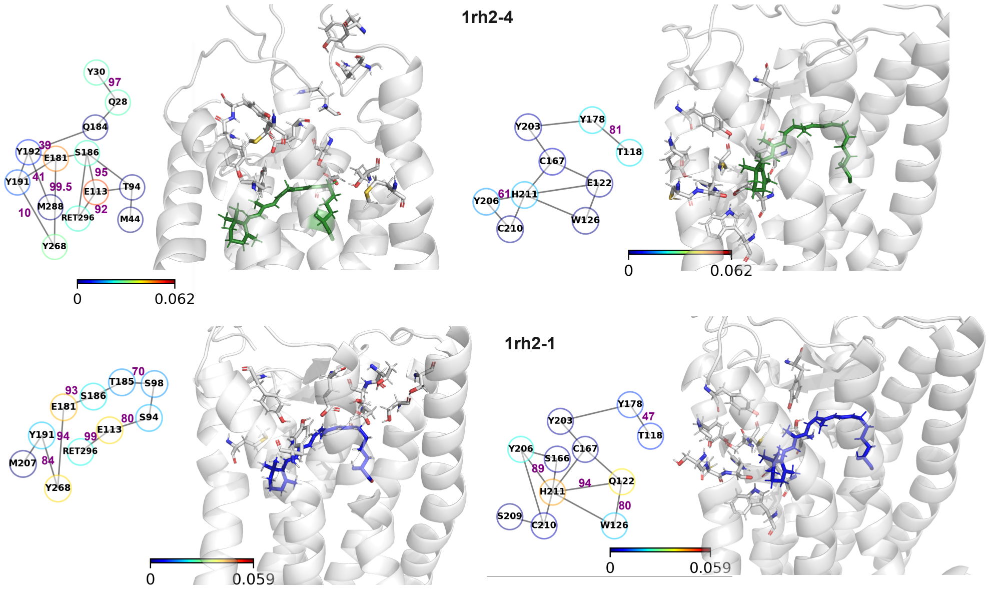


Figure S9: Non-water mediated H-Bond network involving RET(left) and residue 122 (right). The graph of H-bonds identified all nodes that can be reached from RET or 122 via H-bonds. Occupancies are shown in purple. Circle colors represent the normalized degree of centrality, the number of direct H-bonds of that protein group. HBs between $SBP^+$ and E113 are more frequent in 1rh2 than in 4rh2. Bridge2 software.

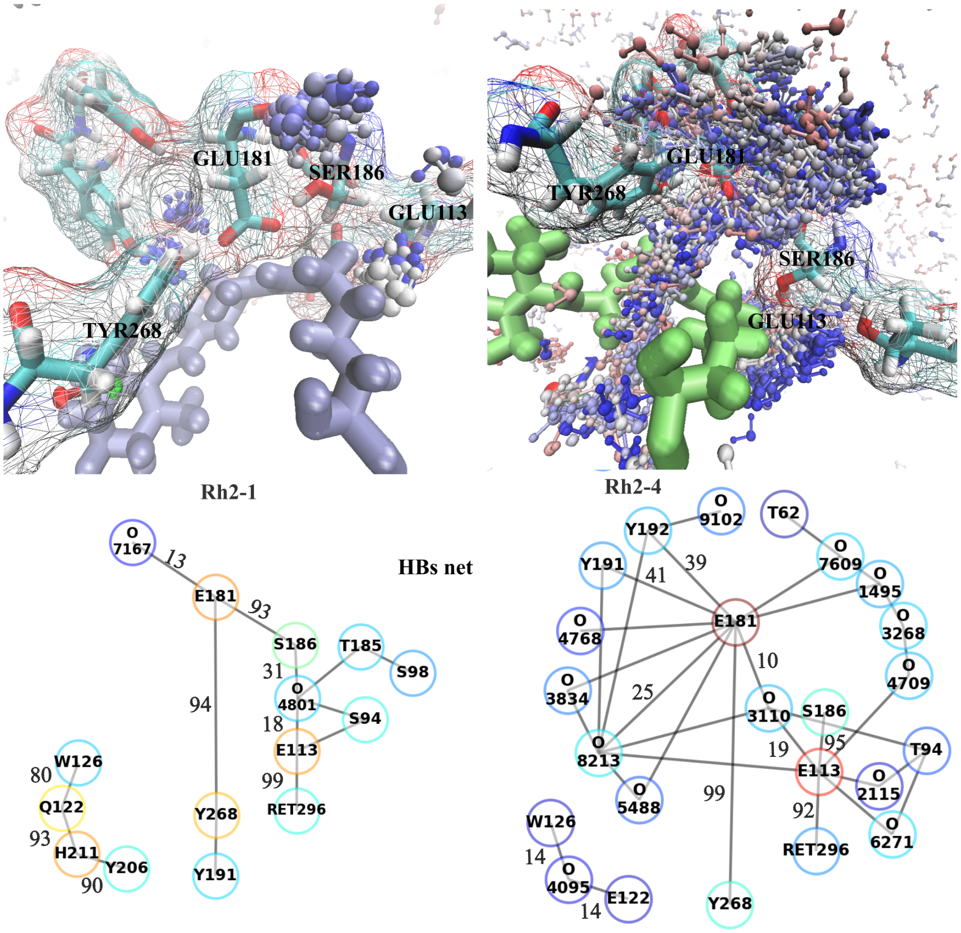


Figure S10: Top) Waters that visit the region near the pocket during MD. The coloring method is trajectory timestep, red means first frames and blue last frames. Blue-sensitive opsin (left) and green-sensitive opsin (right). Bottom) Water-mediated H-Bond network involving RET and residues 122, 113, and 181. The graph of H-bonds identified all interactions of more than 10 % of occupancy during the trajectory via H-bonds. Ocuppancies are shown in black. Circle colors represent the normalized degree of centrality, the number of direct H-bonds of that protein group (the more red the more central)

### References


(1) Corpet, F. Multiple sequence alignment with hierarchical clustering. *Nucleic acids research* **1988**, *16*, 10881–10890.

(2) Chinen, A.; Matsumoto, Y.; Kawamura, S. Reconstitution of ancestral green visual pigments of zebrafish and molecular mechanism of their spectral differentiation. *Molecular*

*biology and evolution* **2005**, *22*, 1001–1010.
